# Isoform-specific regulation of SMARCAD1 dosage in ectodermal homeostasis and dysplasia

**DOI:** 10.64898/2026.08.17.745107

**Authors:** Markus Seibert, Lea Kursawe, Manuel Lang, Emily N. Pöschel, Falk K. Thiemig, Paul Kim, Lea Gutenkunst, Andreas Schlundt, Ritva Tikkanen, Jacqueline E. Mermoud

**Affiliations:** Institute of Molecular Biology and Tumour Research, Philipps University Marburg, Marburg 35043, Germany; Center for Human Genetics, Philipps-University Marburg, Marburg, Germany; University of Greifswald, Institute of Biochemistry, 17489 Greifswald, Germany; Institute of Biochemistry, Medical Faculty, University of Giessen, Friedrichstrasse 24, 35392 Giessen, Germany

**Keywords:** ATP-dependent chromatin remodeller, haploinsufficiency, splicing, UTR, NLS

## Abstract

Haploinsufficiency is a common cause of developmental disorders, yet the mechanisms regulating protein abundance of dosage-sensitive genes remain incompletely understood. SMARCAD syndrome comprises inherited ectodermal dysplasias caused by mutations affecting SMARCAD1-s, the skin-specific short isoform of the ATP-dependent chromatin remodeller SMARCAD1. Here, we show that independent patient-derived mutations impair splicing of this isoform, resulting in intron retention. Although total mRNA abundance is reduced, spliced transcripts remain translationally competent and yield variably reduced levels of wild-type protein, providing evidence for a threshold-dependent haploinsufficiency model of disease. Under physiological conditions, the isoform-specific non-coding exon 1 dampens protein production without altering transcript levels, revealing previously unrecognized post-transcriptional control of SMARCAD1-s dosage. Our findings implicate upstream open reading frames as candidate mediators of translational control. Unexpectedly, SMARCAD1-s exhibits regulated nucleocytoplasmic distribution. Functional analyses identify two nuclear localization signals that differentially contribute to localization of the SMARCAD1 isoforms. Together, our findings clarify the molecular basis of SMARCAD syndrome and identify post-transcriptional and spatial mechanisms controlling the abundance and localization of a dosage-sensitive chromatin remodeller, providing insight into protein dosage control.

## Introduction

Organization of DNA into chromatin is the fundamental stuctural platform which underpins the ability of a cell to safeguard and regulate its genome. The modulation of chromatin structure in response to cellular demands requires ATP-dependent chromatin remodellers. These enzymes shape the packing of DNA by altering nucleosome arrangements and composition, thereby influencing the accessibility of DNA to various factors and regulating fundamental genome activities like transcription, DNA replication and DNA repair ^1,2^. Genetic mutations in chromatin remodelers often result in impairment of embryonic development and are hallmarks of many human diseases, notably various neurological disorders and cancer ^2–4^. More than 20% of human cancers harbor mutations in genes encoding members of the SWI/SNF remodeller family. Recent functional and structural studies have advanced our understanding of the pathogenic consequences, which arise from disrupted chromatin accessibility and mis-regulated transcriptional programs ^4–7^. By contrast, much less is known about the contribution of chromatin remodellers to inherited disorders that result from developmental defects.

Heterozygous mutation in the chromatin remodeller SMARCAD1 (SWI/SNF-related, matrix-associated actin-dependent regulator of chromatin, subfamily A, containing DEAD/H box 1) cause autosomal dominant ectodermal dysplasias (ED), namely Isolated Adermatoglyphia (OMIM, 136000), Basan- (OMIM 129200) and Huriez syndrome (OMIM 181600) ^8–15,15–20^.

A common feature of these rare diseases is the congenital absence of fingerprints due to lack of dermatoglyphs on hands and feet^8,9,21^. This phenotype affects gripping, sweating, thermoregulation, as well as biometric authentication ^21^. Additional clinical manifestations present themselves in Basan and Huriez syndromes, with ∼15% of Huriez patients developing aggressive squamous cell carcinoma with high metastatic potential ^12–20,22^. It was suggested that all three diseases represent the phenotypic spectrum of the same disorder coined SMARCAD syndrome for **S**MARCAD1-associated congenital facial **M**ilia, **A**dermatoglyphia, **R**educed sweating, **C**ontractures, **A**cral Bullae, and **D**ystrophy of nails ^12–15^. Overall, these findings reveal that SMARCAD1 is vital for the formation of ectodermal structures, yet the role of SMARCAD1 in these ED or in normal skin development remains unknown.

Recent work has shown that SMARCAD1 functions as a chromatin remodeller in vitro with a preference for nucleosome intermediates thought to occur during replication fork progression, transcription and repair ^23,24^. Evidence that SMARCAD1 is required during embryogenesis came initially from studies in mice, as its deletion results in developmental abnormalities and prenatal-perinatal lethality ^25,26^. Its presence is required for embryonic stem cell homeostasis and transcriptional silencing of endogenous retroviruses ^25–31^. In humans, two major SMARCAD1 isoforms have been identified, encoding proteins of 1,026 and 596 amino acids, the latter corresponding to the C-terminal portion of the full-length protein. Hereafter, we will refer to these isoforms as long, SMARCAD1-l, and short, SMARCAD1-s, respectively. Both the full-length protein and its truncated form retain the ATPase and helicase characteristic of SNF2-family chromatin remodeller enzymes. Whether these isoforms have distinct or overlapping functions, however, remains unresolved. Studies to date have primarily focused on the ubiquitously expressed SMARCAD1-l isoform and its roles in genome stability, DNA repair, and chromatin organization ^32–38^. However, pioneering studies revealed that SMARCAD syndrome arises specifically from mutations in SMARCAD1-s ^8^ .

SMARCAD1-s contains an alternative first exon (378nt) upstream of the translational start site which resides 9 nucleotides into exon 2 ^8^. Although the function of this non-coding exon 1 is unknown, sequencing of DNA from over 35 patients revealed that all mutations, ranging from individual single nucleotide changes to deletions, cluster around the donor splice site of exon 1 (Figure 1a) ^8–13,13–17,39^. It was subsequently shown that the G>T substitution at the ultra-conserved first nucleotide of the intron causes aberrant pre-mRNA splicing and complete loss of wild-type transcripts in a range of cell types ^8^. This led to the suggestion that disruption of SMARCAD1-s pre-mRNA splicing underlies SMARCAD-syndrome by preventing mRNA maturation and thereby causing a loss-of-function effect ^8,17^. However, the generality of this proposed mechanism was challenged by the observation that a mutation at position 3 of the donor splice site (c.378 +3A>T) exhibited no detectable splicing defect ^16^. These findings raised the possibility that multiple mechanisms contribute to disease pathogenesis and prompted the question of how exactly pathogenic mutations affect SMARCAD1-s enzyme function.

**Figure 1:**
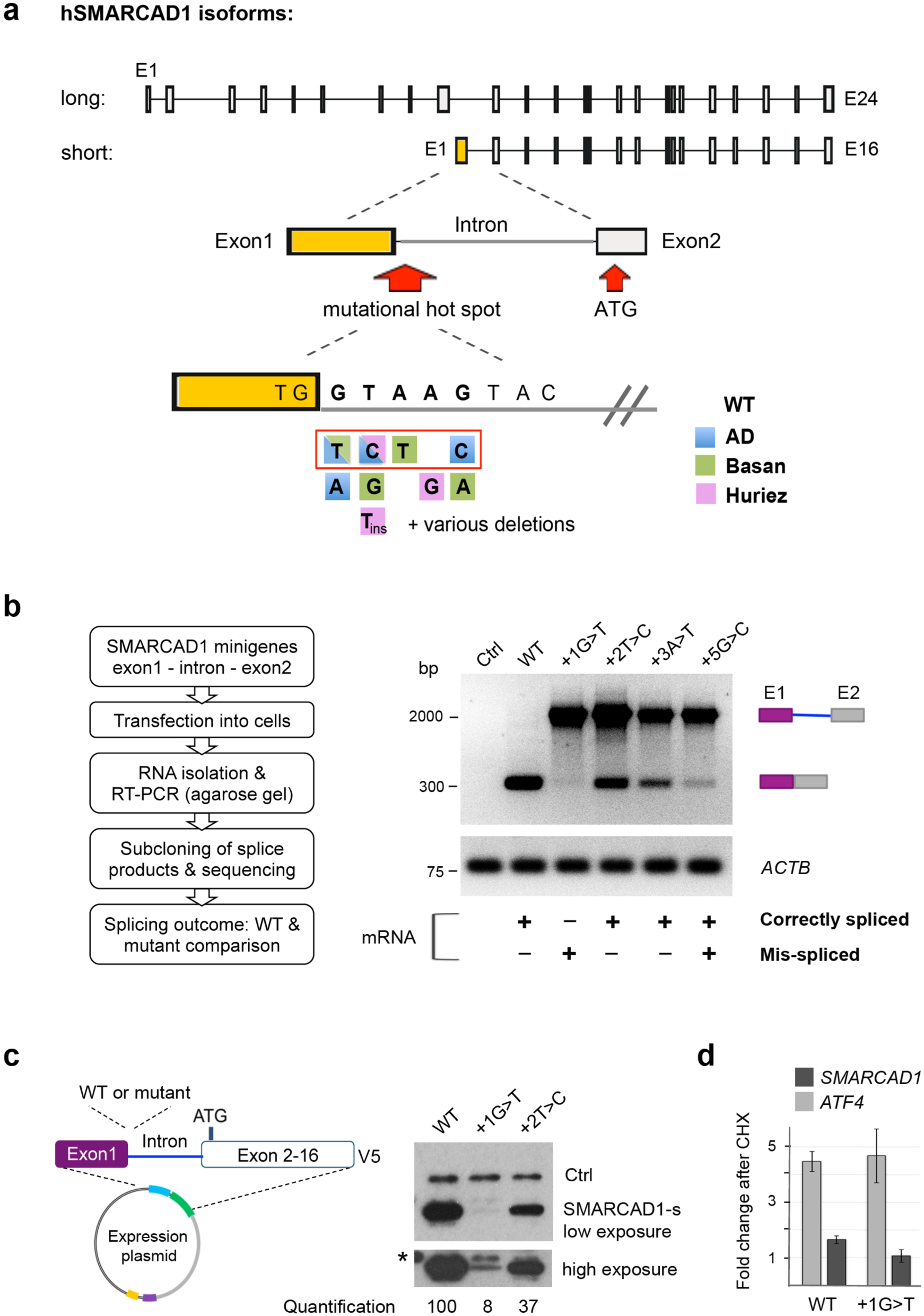
Disease-associated SMARCAD1-s mutations impair splicing and reduce protein abundance. **(a)** Schematic of the two major h *SMARCAD1* isoforms (Chr4q23). The short isoform contains an alternative non-coding exon 1 (yellow) with translation starting in exon 2 (ATG). Point mutations associated with Adermatoglyphia (blue), Basan (green) or Huriez (pink) Syndromes cluster at the exon 1 splice donor ^8–10,12,14–17,19,39^. Deletions of 1 bp to over 1 kb also affect this hotspot and cause clinical manifestations ^10,12,13,17^. Mutations analysed here are boxed in red. **(b)** Mutations alter splicing efficiency but still produce correctly spliced mRNA, except c.378 +1G>T. Left, workflow. Right, representative RT-PCR of minigene transcript (Supplemental Figure 1b) resolved by agarose gel electrophoresis. Pre-mRNA comprising exon 1, intron 1 and exon 2 (2226 bp) and mRNA (355 bp) are indicated. *ACTB,* reference gene; Ctrl, untransfected cells. Sequencing summary shows correctly and aberrantly spliced mRNAs, as detailed in Supplemental Figure 1c. HeLa cells, n=2 biological repeats. **(c)** Short SMARCAD1 protein is produced despite splice site mutations. Expression-plasmids containing all 16 exons of *SMARCAD1*-s, with or without splice site mutations, were transfected and whole-cell lysates analysed by V5 immunoblot to detect the C-terminal tag. The upper band in c.378 +1G>T (asterisk) is non-specific. GFP co-transfection monitored transfection efficiency (Ctrl). Quantification (%) of SMARCAD1-s relative to GFP is shown (V5/GFP). HeLa results (n=2) were reproduced in keratinocytes (Supplemental Figure 1f). **(d)** RT-qPCR of HeLa cells depleted of endogenous SMARCAD1 but transfected with SMARCAD1-s constructs of panel c, WT or c.378 +1G>T, after cycloheximide (CHX, 50 µg/ml) or vehicle treatment for 5 hours. Fold enrichment of *SMARCAD1-s* in CHX over control (±S.D. normalised to *ACTB, GAPDH*) is shown. *ATF4* control is upregulated 4-5 times upon CHX. Technical triplicates are shown; similar results were obtained after 6 hours.

By systematically assessing the consequences of pathogenic mutations on SMARCAD1-s splicing and enzyme production, we show that disease causing variants reduce splicing efficiency but do not abolish synthesis of wild-type protein. SMARCAD1-s protein abundance is variably reduced, supporting a dosage-dependent model of pathogenesis in which disease results from failure to maintain enzyme levels within a functional range. We further establish that upstream non-coding sequences constitute a critical post-transcriptional regulatory layer controlling SMARCAD1-s abundance. Notably, and unusually for a chromatin remodeler, we find that SMARCAD1-s exhibits a nucleocytoplasmic steady-state distribution. We identify two nuclear localization signals in SMARCAD1 and demonstrate that they govern the nuclear entry of the individual isoforms. Taken together, our findings define new principles of isoform-specific regulation and dosage control in chromatin remodellers.

## Results

### Disease linked mutations in SMARCAD1 compromise pre-mRNA splicing

We set out to elucidate whether mutations located at different positions of the donor splice site of exon 1 in SMARCAD1-s disrupt the normal patterns of pre-mRNA splicing and cause loss of SMARCAD1-s protein. Of the mutations described to date, we selected to pursue four (Figure 1a) based on the following criteria: (i) We prioritized base exchanges common to the distinct syndromes. In addition to c.378 +1G>T, reported in both Adermatoglyphia and Basan syndrome ^8,20^, we included c.378 +2T>C, identified in patients with Basan and Huriez syndromes ^9,17^. Positions +1 and +2, are of particular importance, as they constitute the highly conserved dinucleotide at the 5’ end of introns in canonical splice donor sites, and pathogenic variants at these residues are frequent in rare diseases ^40^. (ii) Distance from the core splice site: To evaluate whether base changes further away from the ultra-conserved donor site are tolerant or deleterious for splicing, we also analysed changes at positions 3 (c.378 +3A>T) ^16^ and 5 (c.378 +5G>C) ^9^. (iii) Mutation frequency: More than 40 patients with the selected mutations have been reported ^8,9,17,20,39^. (iv) To explore the contribution of the intrinsic strength of the donor splice site, we selected substitutions representing a range of predicted splice site strengths, as determined by MaxEntScan ^41^. Notably, the c.378 +1G>T and c.378 +2T>C substitutions are predicted to markedly weaken the splice donor site relative to the wild-type sequence (Supplemental Figure 1a).

To directly compare the splicing effects of these mutations within the same assay, we employed a previously validated minigene system comprising exon 1, intron 1 and exon 2 of short SMARCAD1-s ^8^. Single base changes representing the selected donor splice site mutations were introduced into the minigene (Supplemental Figure 1b). As splicing accuracy can vary across different cell types ^42^ we performed the analysis in two backgrounds, cervical carcinoma cells (HeLa’s) and skin cells (HaCaT keratinocytes). The wild-type SMARCAD1-s minigene produced abundant mRNAs species, that we confirmed to be correctly spliced by Sanger sequencing, whereas the corresponding pre-mRNA was barely detectable (Figure 1b and Supplemental Figure 1c, e). Having created conditions that allow very efficient splicing reactions, we then compared the splicing patterns of the WT construct with those carrying mutations at positions +1 up to +5 of intron 1. Remarkably, all mutations tested led to a pronounced accumulation of unspliced transcript compared to the spliced product (Figure 1b and Supplemental Fig 1e), demonstrating that substitutions both at the core donor splice site (+1, +2) and at more distal positions (+3, +5) substantially impair SMARCAD1-s splicing in vitro. This is in contrast to a previous study suggesting that the c.378 +3A>T mutation did not affect the minigene splicing efficiency ^16^. This discrepancy could reflect the cell source, as COS7 (African Green Monkey) kidney cells were used previously, while we employed HeLa cells and human keratinocytes, the later representing the most appropriate system, as confirmed in our expression analysis below.

We conclude that of all four pathogenic mutations lead to increased intron retention in vitro. This results in reduced levels of mature SMARCAD1-s mRNA, although to different extents. Higher levels of spliced products were detected for the c.378 +2T>C and c.378 +3A>T mutation, whereas the c.378 +1G>T and c.378 +5G>C mutations produced markedly lower levels of spliced mRNA (Figure 1b and Supplemental Figure 1e). To investigate whether the detected mRNAs species were accurately spliced, we purified cDNAs and performed Sanger sequencing.

For the c.378 +1G>T substitution, we detected only aberrantly spliced variants in both HeLa and HaCaT cells (Supplemental Figure 1c, e). The mis-splicing events observed are in excellent agreement with previously reported alterations, where transcripts either lacked the terminal G of exon1 (ΔG), or retained an extra 51 bp from intron 1 ^8^. Additionally, we identified another mis-spliced product, lacking 214 bp of Exon 1 but retaining 163 bp of the intron 1. A complete overview of all transcript isoforms is provided in Supplemental Figure 1c and e, with the ΔG variant representing the most abundant species. This is likely due to the fact that the spliceosome recognizes the GT as the boundary of the intron in most genes, and altering the TG/GT wild-type splice site to TG/TT generates a new donor splice (T/GTT).

Importantly, in the context of all splice site mutations studied, the absence of correctly spliced transcripts was the exception in both cell types. In contrast to the c.378 +1G>T variant, we detected correctly spliced mRNA in all other splice site mutants analysed. The c.378 +5G>C substitution yielded mainly correctly spliced products, as well as two distinct mis-spliced variants in HeLa cells (Supplemental Figure 1c, e). Notably, for the c.378 +2T>C and c.378 +3A>T mutations we detected exclusively correctly spliced mRNA (Supplemental Figure 1c, e).

Overall, the splice site mutants fall into two groups. Group 1 comprises mutations that cause predominantly aberrant splicing, with little or no correctly spliced mRNA (c.378 +1G>T). Group 2 comprises mutations where correct splicing does occur, albeit at a consistently lower level than in the WT (c.378 +2T>C, c.378 +3A>T, c.378 +5G>C).

### Splicing defects do not completely abolish protein translation of short SMARCAD1

Our finding that mRNA could still be detected despite mutations in the donor splice site of exon 1 prompted us to investigate whether these transcripts produce protein. We generated constructs that drive the expression of all 16 exons of short SMARCAD1 with a V5 tag at the C-terminus to allow detection of full-length proteins. These constructs all retained exon1 and intron 1 sequences (Figure 1c, left). We compared the WT with representative splice site mutations from Group 1, which display aberrant splicing (c.378 +1G>T), and Group 2, which exhibit predominantly correct splicing. The c.378 +2T>C and c.378 +1G>T substitutions were selected as they represent the most frequent mutations found in patients ^8,9,17,20^. Constructs were introduced into HeLa cells together with a second plasmid expressing a GFP-fusion protein to ensure comparable transfection efficiency and to evaluate the relative levels of SMARCAD1-s (Supplemental Figure 1d, workflow; Figure 1c, right, lane 1).

The c.378 +2T>C mutation did not abolish SMARCAD1-s protein production, as significant protein levels were detected in both Hela cells and keratinocytes (Figure 1c, right, lane 3 and Supplemental Figure1f). While the levels of protein are lower than when the splice site was wild type, this result is consistent with the RT-PCR results, which revealed a relative high proportion of unprocessed transcripts (Figure 1b, Supplemental Figure 1e). Together, these observations indicate that reduced mRNA levels result in reduced protein levels, suggesting that those transcripts that are correctly spliced proceed to translation.

Next, we examined the fate of transcripts that are aberrantly spliced and determined whether they retain translational competence. We tested whether SMARCAD1-s protein can also be generated from the c.378 +1G>T mutant construct, for which neither we nor previous investigators detected correctly processed mRNA (Figure 1b and Supplemental Figure 1c, e) ^8^. In this setting, SMARCAD1-s protein levels were markedly reduced. Immunoblot analysis revealed a substantial reduction to less that 10% of the WT protein (Figure 1c and Supplemental Figure 1f). Consistent with this, the c.378 +1G>T mutant displayed even lower mRNA levels than the c.378 +2T>C variant (Figure 1b and Supplemental Figure 1e). Treatment of cells with cycloheximide (CHX) to block mRNA de-capping and translation elongation (Schwarz and Parker, 1999; Mercier et al 2024; Park et al 2017), resulted in RNA stabilization of a control transcript (Figure 1d, ATF4). In contrast, the c.378 +1G>T mutant minigene gave rise to RNAs that are not affected by CHX, suggesting reduced splicing efficiency as the major cause of low mRNA levels.

To further explore this, we analysed a construct designed to mimic the predominant splicing outcome of the c.378 +1G>T mutant, namely the deletion of the terminal G of exon1 (ΔG) (Figure 2a left). This construct recapitulates an aberrant, yet fully efficient, splicing event that uses a cryptic splice site. If such mis-spliced transcripts were non-functional and subject to elimination, no protein would be expected. However, the ΔG mutant produced SMARCAD1-s protein levels comparable to the-control construct containing a WT intron 1 (Figure 2a,

**Figure 2:**
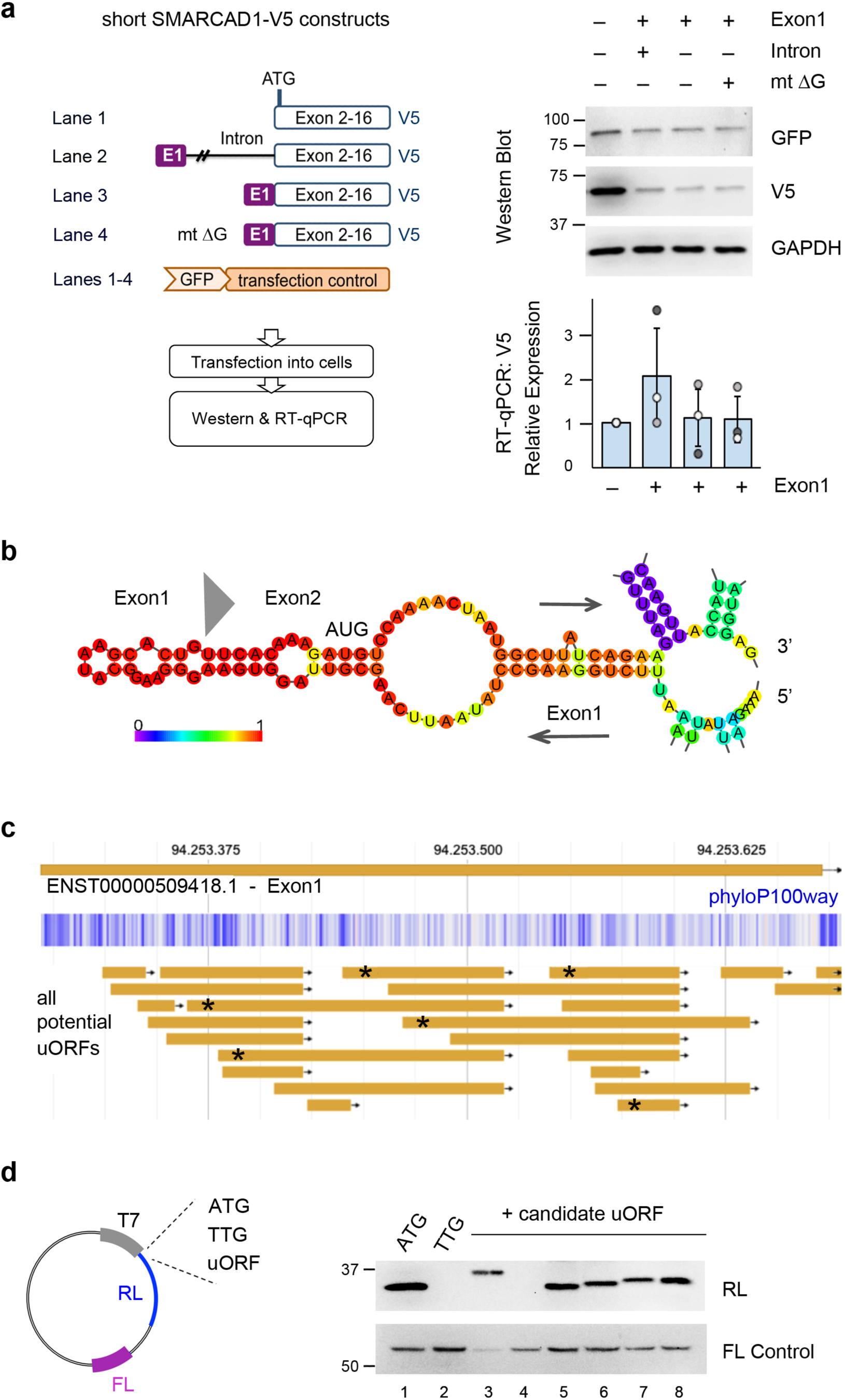
The SMARCAD1-s-specific non-coding exon 1 post-transcriptionally regulates protein abundance. **(a)** Exon 1 reduces short SMARCAD1 protein levels without affecting RNA. Left, workflow. HeLa cells were transfected with different *SMARCAD1-s* constructs, containing the same promotor, *SMARCAD1*-*s* Kozak sequence and C-terminal V5-tag. A co-transfected GFP vector served as control and analysis was carried out 24 hours after transfection. Right, top: Representative Western blot (n=3-6) showing V5 (SMARCAD1-s), GFP (transfection control), and GAPDH (loading control). Right, bottom: RT-qPCR of V5 mRNA with three individual biological replicates shown as dots, normalized to Actin, GAPDH and GFP. Errorbars represent the mean ± S.D., n = 3. No significant differences were observed in a two-tailed Student’s *t*-test. **(b)** RNA secondary structure of *SMARCAD1-s* predicted with RNAfold (ViennaRNA Package) reveals a high propensity for the AUG region in exon 2 to be stably structured in the presence of exon 1. Heat bar indicates base-positional probability. Translation start codon and exon1- 2 junction are marked in this enlarged segment of the full predicted exon 1/ 2 structure shown in Supplemental Figure 2b. **(c)** Exon 1 of SMARCAD1-s transcript contains more than 20 candidate upstream open reading frames (uORFs), annotated via Ribo-uORF ^87^. Visualization in JBrowse corresponds to chr4:94.253.294-94.253.671. Candidate uORFs are defined as start (ATG/CTG/GTG/TTG/ACG)-to-stop (TAG/TGA/TAA) with a minimum length of 18 nt. Blue tracks indicate evolutionary conservation across the exon 1 sequence (PhyloP100way). **(d)** Translation from putative uORFs in exon 1 of SMARCAD1-s assessed by in vitro cell-free translation. Western blot detecting Renilla Luciferase (RL) from a psi-CHECK2 Reporter which contains a T7 promoter (lane 1). The ATG start codon of the RL gene was mutagenized (lane 2) and replaced with putative uORFs (lanes 3-8) to assess their ability to initiate translation in vitro. Firefly Luciferase (FL) served as control. Each construct was tested at least twice.

Western blot, compare lanes 2 and 4). Collectively, these findings indicate that the c.378 +1G>T mutation results in severely diminished, but not entirely attenuated, production of SMARCAD-1-s protein.

We conclude that 5’ splice site mutations in intron 1 of SMARCAD1-s do not necessary impair protein production from downstream exon 2 per se, but reduced mRNA abundance correlates closely with diminished protein levels. This suggests that, in patients carrying such mutations, the absolute amount of SMARCAD1-s protein is critical for normal function, consistent with current models of haploinsufficiency.

### Exon1 is critical for tuning SMARCAD1-s protein levels

To understand how lowering the levels of SMARCAD1-s activity contributes to disease, it is necessary to understand the normal expression, regulation and subcellular localization of SMARCAD1-s. First, we examined the critical parameters that determine the abundance of SMARCAD1-s protein. Several studies have demonstrated that introns can modulate gene expression by affecting nuclear export, transcript stability or translation efficiency ^43^. In some cases, these effects are mediated by intronic sequences per se, whereas in others they depend on the process of splicing. We found that deletion of intron 1 had only a minor impact on SMARCAD1-s protein levels compared with expression constructs that retain the intron (Figure 2a, Western, compare lane 2 and lane 3). The latter harbors the first 1, 358 bp and the last 500 bp of the 11,035 bp long intron1 ^8^. We infer that neither the intron1 sequences flanking the donor and acceptor splice sites nor splicing per se play major roles in determining SMARCAD1-s protein amounts.

Next, we considered whether SMARCAD1-s protein abundance is affected by protein stability. One of the major pathways regulating protein turnover is the ubiquitin proteasome system, which can be blocked by the compound MG132. Inhibitor treatment effectively prevented degradation of known proteasome targets such as p53, but had no detectable effect on either SMARCAD1 isoforms (Supplemental Figure 2a). These findings argue against a key role of the proteasome pathway in the control of steady-state SMARCAD1-s protein levels.

Untranslated regions (UTRs) of mRNAs typically play key roles in modulating the abundance of the corresponding protein ^44^. To test whether exon 1, which comprises 363 bp of non-coding sequence, affects SMARCAD1-s expression, we deleted it from SMARCAD1-s expression-constructs. Strikingly, removal of exon 1 resulted in a marked increase in SMARCAD1-s protein levels compared to constructs containing exon 1 (Figure 2a, Western blot, compare lane1 with 2). These results show that exon 1 plays a major role in regulating SMARCAD1 protein abundance.

Two main mechanisms could explain this observation: exon 1 might impact SMARCAD1-s mRNA stability or translation efficiency. Although protein levels were substantially reduced in the presence of exon 1, this difference was not reflected in RNA levels, as RT-qPCR showed no significant increase in SMARCAD1-s transcripts in samples lacking exon 1 (Figure 2a, right panel, bottom, compare bar 1 with 2-4). We therefore conclude that reduced protein levels are not due to decreased mRNA abundance.

To explore whether exon 1 modulates translation, we performed computational RNA-secondary structure analyses. Most eukaryotic mRNAs are translated by a scanning mechanism in which the ribosomal pre-initiation complex migrates along the transcript, locally unwinding RNA structures until encountering a suitable translation start codon. RNA folding predictions indicated that inclusion of exon 1 increases the structural stability around the AUG start codon in exon 2 (Figure 2b and Supplemental Figure 2b). This supports a model in which exon 1 reduces translation efficiency by restricting ribosomal access to the AUG, thereby dampening SMARCAD1-s protein synthesis.

Well-established examples of principal cis-acting elements that regulate translation are upstream open reading frames (uORFs), that occur within untranslated regions of almost half of all human transcripts ^45,46^. Their translation typically attenuates expression of the main ORF ^47^, which resembles the phenotype that we observe here. Therefore, we examined exon 1 for potential functional uORFs, defined by a start codon, comprising an AUG or a non-canonical codon, and a corresponding in-frame stop codon located upstream of the coding sequence. We identified more than 20 candidate uORFs in exon 1 by bioinformatic prediction (Figure 2c). This number exceeds what would be expected by chance, given that uORFs are generally underrepresented in the 5′ UTRs of eukaryotic mRNAs ^45^ and in line with the observation that developmental and dosage-sensitive genes typically harbor more uORFs than genes tolerant to loss of function ^47,64,65^. Phylogenetic analysis (phyloP) revealed that a subset of these putative uORFs possess evolutionary conserved start codons (Figure 2c, conserved sites are shown in blue), suggesting a selective pressure to maintain these defining features, likely due to functional constraints ^45^. We selected six candidate uORFs, covering the length of exon 1 and encompassing a range of sizes from 3-55 amino acids (Figure 2C, marked with an asterix), and tested their ability to restore translation to a reporter gene lacking a functional AUG (Figure 2d). All but one of the tested uORFs drove the translation of the reporter in a cell free system, indicating that several of the predicted uORFs can indeed function as translation initiation sites, consistent with a model in which uORFs regulate the translation of SMARCAD1-s. When Exon 1 is skipped, all uORFs are eliminated, the 5’ UTR collapses and the ribosome encounters the main ATG almost immediately after the cap. The net effect is likely derepression of translation as we observe in Figure 2a.

In vivo, uORF activity is context-dependent, influenced by cellular state, experimental conditions, and tissue type ^48^. It will therefore be an endeavor for the future to gather experimental support for the possibility that uORFs control endogenous SMARCAD1-s expression. A prerequisite will be the selection of the appropriate cell source and state. This requires the knowledge of where short SMARCAD1 transcripts occur in the body, which is what we set out to answer below.

### SMARCAD1-s is predominantly expressed in ectoderm derived cell lineages and peaks in proliferating keratinocytes

Within the *SMARCAD1* locus, several promoter regions are predicted, including one associated with the short isoform located at chr4:94253114-94253350 (hg38) (Supplemental Figure 3a). This indicates that distinct transcription-start-sites could drive the expression of the different *SMARCAD1* isoforms, likely underpinning tissue-specific gene expression ^49^. As a full understanding of the expression profile of *SMARCAD1-s* and its relationship to the ubiquitously expressed long isoform is lacking, we first examined genome-wide maps of transcriptional start sites assembled from every major human organ, multiple primary cell types and over 200 cancer cell lines by the FANTOM5 consortium (Functional ANnoTation Of the Mammalian genome). Across 1829 samples, *SMARCAD1-s* expression was detected in ∼70 (Supplemental Figure 3b; complete list Supplemental Table 1), most abundantly in skin and fingernails, consistent with prior reports ^8,17^. The high levels present in stratified squamous epithelial tissues, such as the epidermis, tongue and esophagus, support a role for this SMARCAD1 variant in epithelial biology. Moderate expression was also observed in neuronal tissues, particularly in the cerebrum, pointing towards additional functions of SMARCAD1-s in the central nervous system (Supplemental Figure 3b). Notably, *SMARCAD1-s* transcripts were found in both fetal and adult tissues and in several cancers of epithelial origin (Supplemental Figure 3b), indicating developmental persistence and disease-associated activation of the short promoter.

Next, we examined *SMARCAD1* expression patterns in the skin, focusing on keratinocytes and fibroblasts, the main cell types of epidermis and dermis, respectively. By RT-PCR analysis, we detected both isoforms in keratinocytes, whereas the short transcript was undetectable in two fibroblast lines (Supplemental Figure 3c). This is in contrast to a previous tissue cDNA blot which suggested high expression of *SMARCAD1-s* in fibroblasts ^8^. Although fibroblasts vary by layer and location, the FANTOM5 CAGE expression atlas confirmed *SMARCAD1-s* expression only in epidermal keratinocytes but not in dermal or normal skin fibroblasts. Based on these findings, we chose keratinocytes as the model system for subsequent analyses.

We obtained skin biopsies from nine adults and isolated primary keratinocytes (KC) for quantitative PCR analysis with isoform-specific primers to estimate the relative abundance of the two *SMARCAD1* isoforms, which are typically not distinguished in genome-wide or single cell data sets (Figure 3a). The long isoform showed relatively consistent expression across donors, whereas the short isoform exhibited greater variability (Figure 3a, b). Comparison of their expression levels revealed no significant correlation between the two isoforms (Figure 3b, Pearson’s *r*=0.31; *p*=0.42).

**Figure 3.**
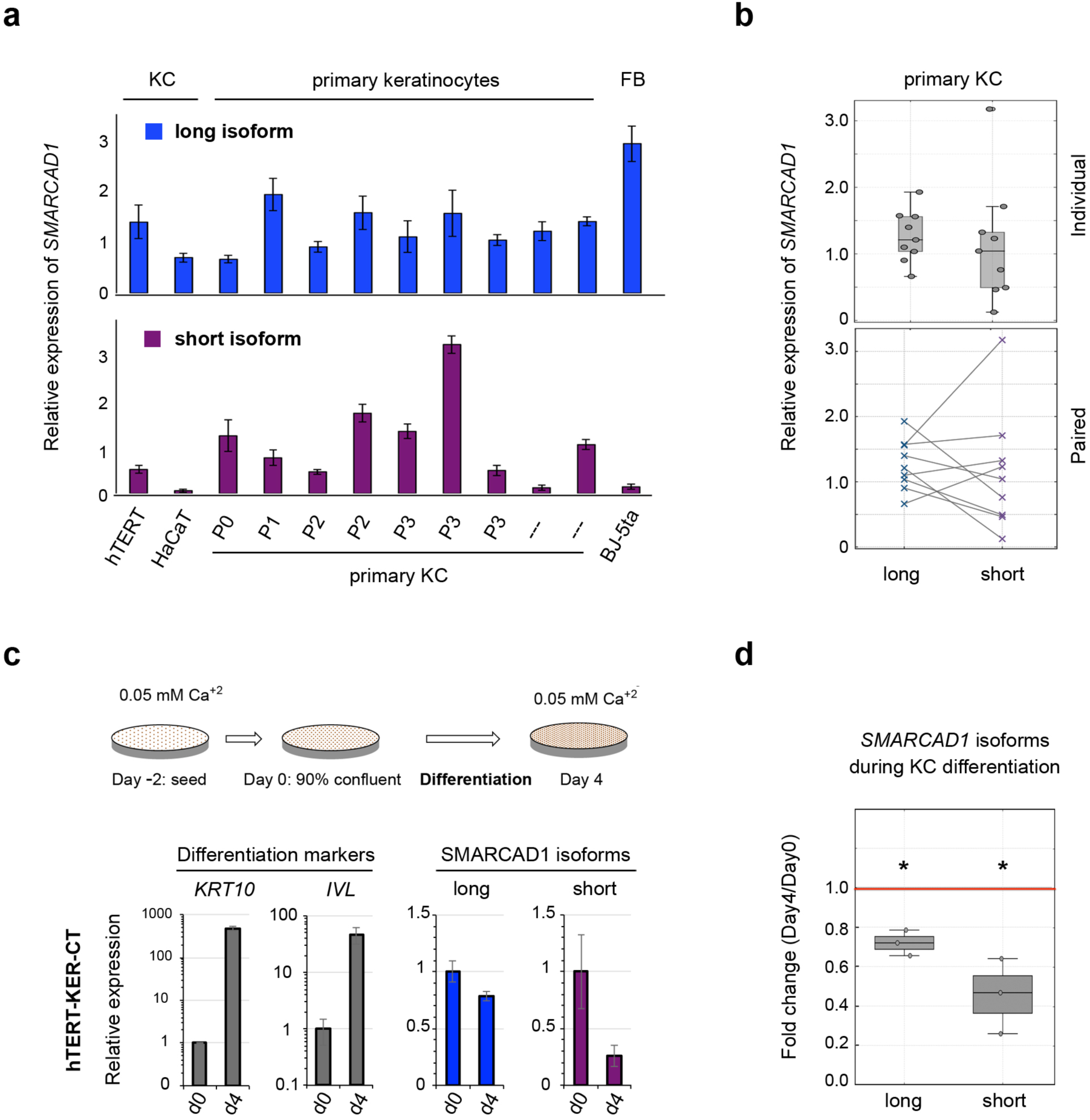
*SMARCAD1* isoform expression in human skin cells. **(a)** RT-qPCR analysis of *SMARCAD1* in keratinocytes (KC: primary, hTERT-KER-KC, HaCaT) and fibroblasts (FB: BJ-5ta). Top, long isoform. Bottom, short isoform. Primary KCs from nine individuals with breast reductions were analysed to assess inter-donor variations. Samples are arranged by passage number (P) where known. Expression is mean ± SD of technical replicates, normalized to *UBC* and *YHWAZ*. Multiple primer pairs (n=2) and independent cDNA preparations (n=1-3) gave consistent results. **(b)** *SMARCAD1* expression in primary KCs from nine donors as depicted in panel a. Top: Isoform specific data from individual donors are shown as circles in a boxplot; median and interquartile range are indicated, with whiskers representing the full expression range. No significant difference between the isoforms (paired *t-*test: *t*= 0.40, *p*= 0.70). Bottom: paired comparison of long versus short isoform per donor; lines connect matched samples from the same donor. Pearson correlation showed no significant correlation between long and short isoform expression (*r*= 0.30, *p*= 0.42). **(c-d)** Downregulation of *SMARCAD1-s* during keratinocyte differentiation. **(c)** hTERT-KER-CT keratinocytes were differentiated by cell-cell contact and harvested at day 0 and day 4. RT-qPCR assessed expression of *SMARCAD1-l* and *SMARCAD1-s*, alongside differentiation makers *KRT10* (keratin 10, early marker) and *IVL* (involucrin, late marker). Expression values were normalized to *YHWAZ* and *UBC* and are represented as mean ± S.D of technical triplicates. **(d)** Box-and-whisker plot shows fold change (day 4/day 0) of *SMARCAD1* isoforms across three independent datasets from panel c and Supplemental Figure 2d, plotted as dots. The central line denotes the median, the box the interquartile range and whiskers extend to the maximum and minimum observed values. Both isoforms were significantly downregulated relative to day 0 (set to 1, red line): *SMARCAD1-l p*= 0.017; *SMARCAD1-s p*= 0.034 (one-sample, two-tailed *t*-test against a hypothetical mean of 1).

In parallel, we evaluated *SMARCAD1* expression patterns in established cell culture models, including spontaneously transformed nontumorigenic keratinocytes from trunk skin (HaCaT) and human telomerase-immortalized foreskin cell lines (hTert-KER-CT keratinocytes; BJ-5ta fibroblasts) (Figure 3a). While *SMARCAD1-l* is expressed highly in BJ-5ta fibroblasts, *SMARCAD1-s* is not, yet it is present in foreskin-derived keratinocytes. We note that *SMARCAD1* transcripts were low in HaCaT cells, consistent with the reported loss or structural alterations of chromosome 4, which harbors the *SMARCAD1* gene ^50^.

As a self-renewing tissue, the epidermis undergoes continuous turnover and differentiation. Closest to the dermis, in the basal layer, reside keratinocytes with a high proliferative potential. They undergo a gradual differentiation program as they migrate upwards towards the outer layer of the skin where they ultimately become cornified cells. In our analysis of primary keratinocytes, we noticed that *SMARCAD1-s* transcript clustered with markers of proliferation rather than differentiation (Supplemental Figure 3d). To directly examine the relationship between *SMARCAD1* expression and keratinocyte differentiation status, we compared isoform expression across two independent differentiation models that recapitulate epidermal differentiation: cell–cell contact inhibition ^51^ and calcium-induced differentiation ^52^. Cells were harvested before and four days after induction of differentiation, and transcript levels were quantified by RT–qPCR (Figure 3c and Supplemental Figure 3e). The analysis was performed in transformed and primary keratinocytes, and expression of canonical epidermal markers confirmed the progression of differentiation. Across all systems, both *SMARCAD1* isoforms were abundantly expressed in undifferentiated keratinocytes and were consistently downregulated upon differentiation. Statistical analysis confirmed that this reduction was significant for both isoforms (Figure 3d; *SMARCAD1-s*: *p* = 0.034; *SMARCAD1-l*: *p* = 0.017). These findings indicate that the short *SMARCAD1* isoform is highly expressed in proliferating keratinocytes, consistent with a role in early epidermal development and dermatoglyph formation, which initiates in the basal layer of the epidermis ^53^.

### SMARCAD1-s is located both in the nucleus and cytoplasm

SMARCAD1-s protein comprises amino acids 430 to 1026 of SMARCAD-l, hence, as it is identical to the C-terminal part of SMARCAD1-l, there are no antibodies that exclusively recognize SMARCAD1-s. To distinguish between the two isoforms and to assess their subcellular distribution, we transiently transfected EGFP-tagged versions into primary and immortalized keratinocytes. As anticipated for a chromatin remodeller, SMARCAD1-l displayed strong nuclear accumulation with no significant signal in the cytoplasm (Figure 4a, long). This is in agreement with the localization of this isoform reported in cell types as diverse as human cancer cells, mouse brains or stem cells ^25,32,54,55^. Nuclear localization was further confirmed by indirect immunofluorescence in untransfected keratinocytes using an antibody specific to the N-terminal region of the long isoform (Supplemental Figure 4a), demonstrating that the tagged protein faithfully recapitulates localization of endogenous SMARCAD1.

**Figure 4.**
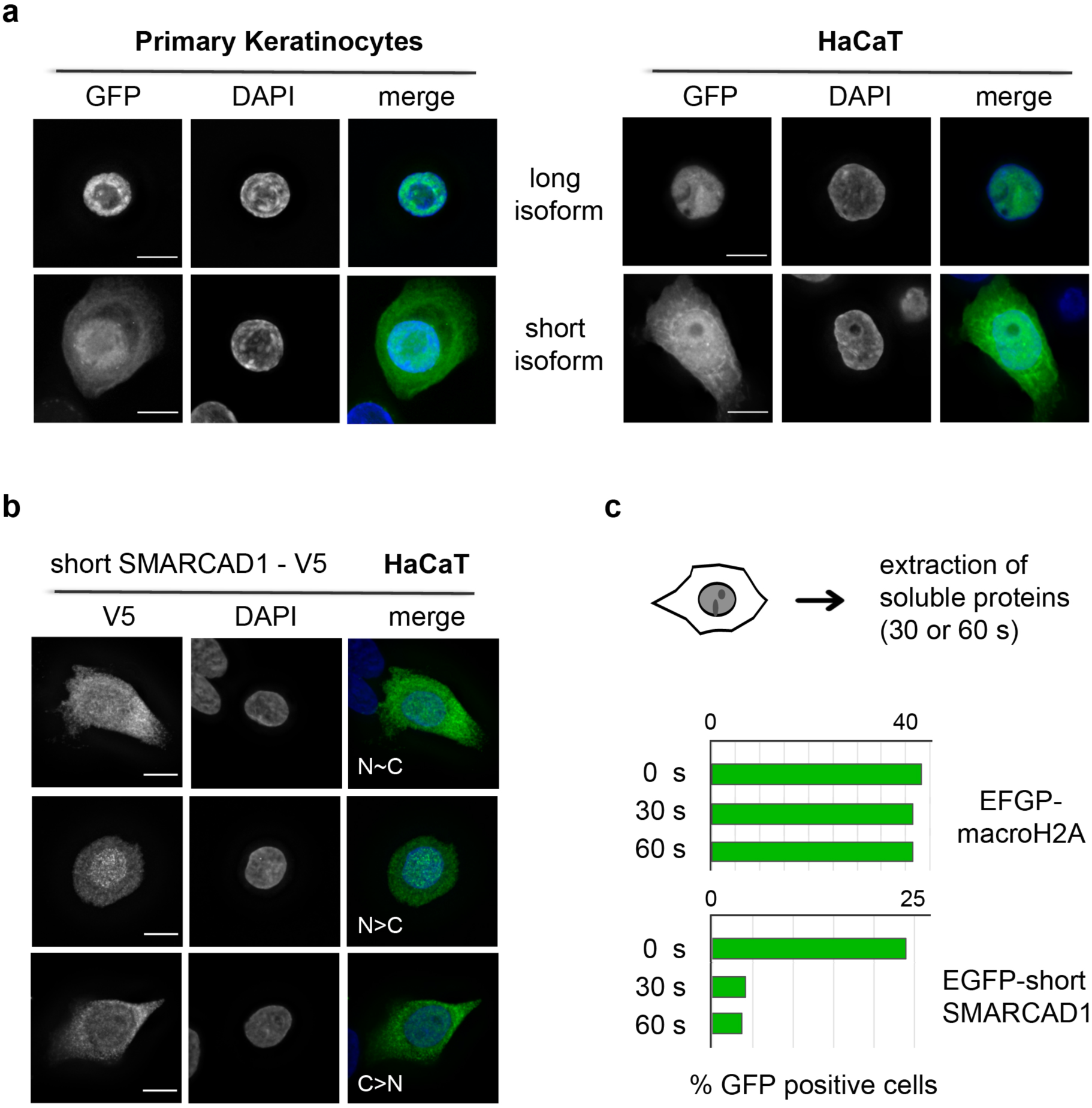
SMARCAD1-s is both nuclear and cytoplasmic and readily extractable. **(a)** EGFP-tagged SMARCAD1 isoforms were expressed in primary (n=3) and immortalized (n=2) keratinocytes (KC), and GFP localization was assessed by fluorescence microscopy. The long isoform was exclusively nuclear and did not redistribute to the cytoplasm upon overexpression (n>100). The short isoform localized to both nucleus and cytoplasm (primary KC: n =116; HaCaT n= 115 GFP-positive cells). **(b)** A C-terminally V5-tagged SMARCAD1-s construct was transiently transfected into HaCaT KCs (n=2). Immunofluorescence with anti-V5 antibody revealed both nuclear (N) and cytoplasmic (C) distribution in all GFP positive cells (n=104). Localization patterns were classified as N∼C (similar nuclear and cytoplasmic signal intensity); N>C (nuclear-enriched) or C>N (cytoplasmic-enriched). **(a-b)** Representative deconvoluted images (Leica DM5500) are shown. Nuclei were counterstained with DAPI. Scale bar, 10 µm. **(c)** Cells expressing EGFP-tagged macroH2A1.2 or SMARCAD1-s were subjected to soluble protein extraction. The proportion of GFP-positive cells was quantified prior to (0 s) and after 30 or 60 seconds of extraction; over 200 DAPI-stained cells were scored per condition. Representative images are provided in the Supplement.

Unexpectedly, EGFP SMARCAD1-s (84kDa) localized to both the nuclear and cytoplasmic compartment, with a robust cytoplasmic accumulation observed in all cells (Figure 4a, short; Supplemental Figure 4b). Importantly, nucleocytoplasmic localization was consistently observed across all transfected cells, irrespective of the EGFP expression levels or amount of DNA input. Moreover, SMARCAD1-s localization was independent of tag position or size, as both C-terminal V5 and N-terminal GFP fusions showed nucleocytoplasmic distribution (Figure 4a, b). In a substantial fraction of keratinocytes, SMARCAD1-s was distributed evenly between the nucleus and cytoplasm (N∼C; >45%), while other cells displayed predominant localization to either compartment (N>C or C>N) (Figure 4a, b; Supplemental Figure 4b). This cell-to-cell variation in protein distribution may reflect dynamic exchange between different compartments, distinct keratinocyte subtypes or differences in cell cycle stage.

In conclusion, SMARCAD1 isoforms exhibit distinct subcellular distributions in skin cells. While tagged SMARCAD1-l is exclusively nuclear, SMARCAD1-s is both nuclear and cytoplasmic. This nucleo-cytoplasmic pattern is consistently observed across all cells, suggesting it represents a steady-state feature of normal cycling keratinocytes. These findings raise the possibility that the function of SMARCAD1-s is not restricted to the nucleus, but that it may also play a role in the cytoplasm.

We next tested if SMARCAD1-s is tightly associated with chromatin or cytoplasmic structures. SMARCAD1-l has previously been shown to be readily extractable from the nucleus, except in S-phase cells, when it engages in DNA replication through protein interactions involving its N-terminal domain, that is absent in SMARCAD1-s ^32,55,56^. We carried out a similar analysis and treated cells with detergent and salt after transient transfection of EGFP-tagged proteins, thereby washing out soluble proteins prior to performing microscopy. In contrast to the chromatin bound control macroH2A1.2, the majority of SMARCAD1-s was readily extractable in both compartments, arguing against it being stably incorporated into chromatin or other large, detergent-resistant complexes (Figure 4c and Supplemental Figure 4c). These findings indicate that SMARCAD1-s is largely soluble and may exert its functions through dynamically regulated interactions.

### Determinants of SMARCAD1-s cellular distribution

Earlier studies in yeast suggested that SMARCAD1-l forms homodimers ^57^, but whether the human long and short isoforms interact remains unknown. To test whether SMARCAD1-l influences the localization of SMARCAD1-s, for example through dimerization, we used HeLa cells, which endogenously express only the long isoform. As in keratinocytes, EGFP-SMARCAD1-s was detected in both the nucleus and cytoplasm (Supplemental Figure 5a). Depletion of SMARCAD1-l had no effect on this distribution. In knockdown cells, SMARCAD1-s maintained identical localization patterns and frequencies (Figure 5a) ^32^. Thus, the subcellular distribution of SMARCAD1-s is independent of SMARCAD1-l.

**Figure 5:**
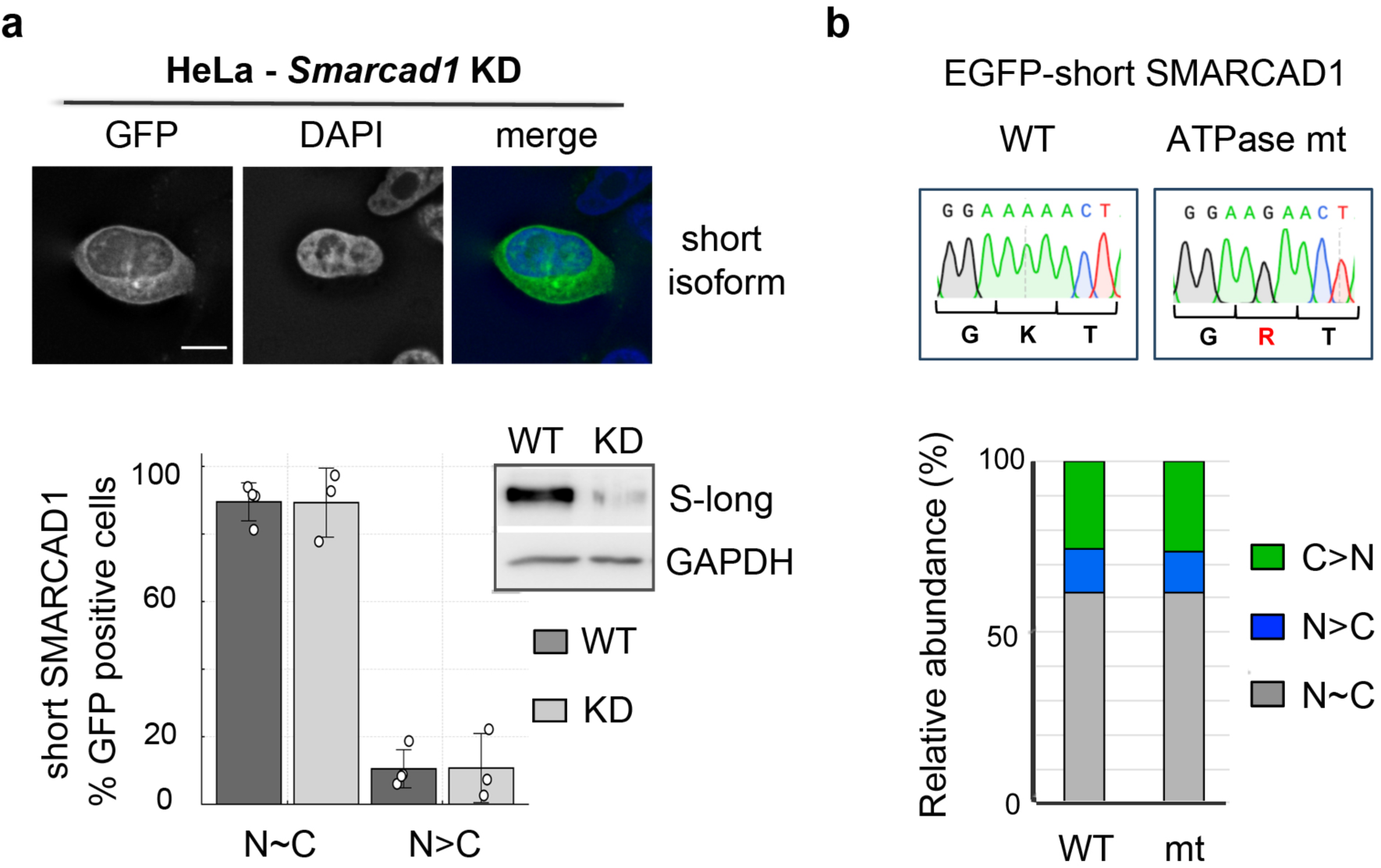
Subcellular localization of SMARCAD1-s is independent of enzymatic activity and SMARCAD1-l. **(a)** Localization of the short, shRNA resistant EGFP-tagged SMARCAD1 isoform was compared in WT and *Smarcad1* knockdown HeLa cells. Representative images are shown on top and in Supplemental Figure 5A; scale bar, 10 µm. Bar graphs show mean ± SD of EGFP-SMARCAD1-s classified as N∼C (similar nuclear and cytoplasmic signal) or N>C (nuclear enriched but cytoplasmic signal present) with dots representing individual data points. No significant difference was detected between WT (n=4; 183 GFP positive cells) and KD (n=3; 115 GFP positive cells) via unpaired two-tailed t-test. Inset: immunoblot confirming SMARCAD1 knockdown; GAPDH, loading control. **(b)** Mutation of the ATPase domain (mt; GKT to GRT) did not alter the localization relative to WT. 125 GFP-positive cells were scored per condition across five slides and categorized as indicated. Representative images are shown in Supplemental Figure 5d.

To assess whether ATPase activity contributes to the localization of SMARCAD1-s, we generated an ATP-hydrolysis-deficient mutant (K98R), previously shown to inactive SWI/SNF ATPases. The localization of this mutant protein was indistinguishable from the wild-type SMARCAD1-s (Figure 5b, quantification; Supplemental Figure 5b, representative images), indicating that the nucleocytoplasmic distribution of the short isoform is uncoupled from its enzymatic activity.

### Distinct nuclear localization signals regulate the distribution of SMARCAD1 isoforms

Nuclear localization of SMARCAD1-l has been attributed to a putative nuclear localization signal (NLS) within a region common to both isoforms ^58^. However, our data show that the short isoform localizes to both the cytoplasm and nucleus, indicating that other sequence elements contribute to the control of subcellular distribution. This could be due to the presence of a cryptic nuclear export signal active in SMARCAD1-s but not in SMARCAD1-l and/or an additional NLS in the amino terminal part of SMARCAD1-l. We therefore next employed bioinformatic analyses to pinpoint putative NLS motifs in SMARCAD1.

Classic nuclear localization signals (cNLS) contain one or two clusters of basic amino acids, referred to as monopartite or bipartite signals, respectively ^59,60^. We used the Protein Subcellular Localization Prediction Tool (PSORT II) ^61^, which relies on cNLS consensus motifs, and the NLStradamus program, which predicts NLS based on stretches of basic residues^62^. The latter identified a monopartite cNLS resembling the SV40 large T-antigen motif (FNKKRKKN) at amino acids 340-347, which we designated NLS1 (Figure 6a and Supplemantal Figure 6a). This motif is absent from the short isoform (Figure 6a). PSORT II highlighted a bipartite cNLS spanning residues 721 to 739 (RRVKEEVLKQLPPKKDRIE), corresponding to the site previously proposed by Adra et al (2000). We refer to this as NLS2 (Figure 6a). Both candidate motifs meet the criteria of a functional NLS: (i) high sequence conservation from mouse to man (Figure 6a), and (ii) predicted surface accessibility in the folded protein. AlphaFold2-based structural modeling indicates that both motifs are exposed in the SMARCAD1 structure (Supplemental Figure 6b).

**Figure 6:**
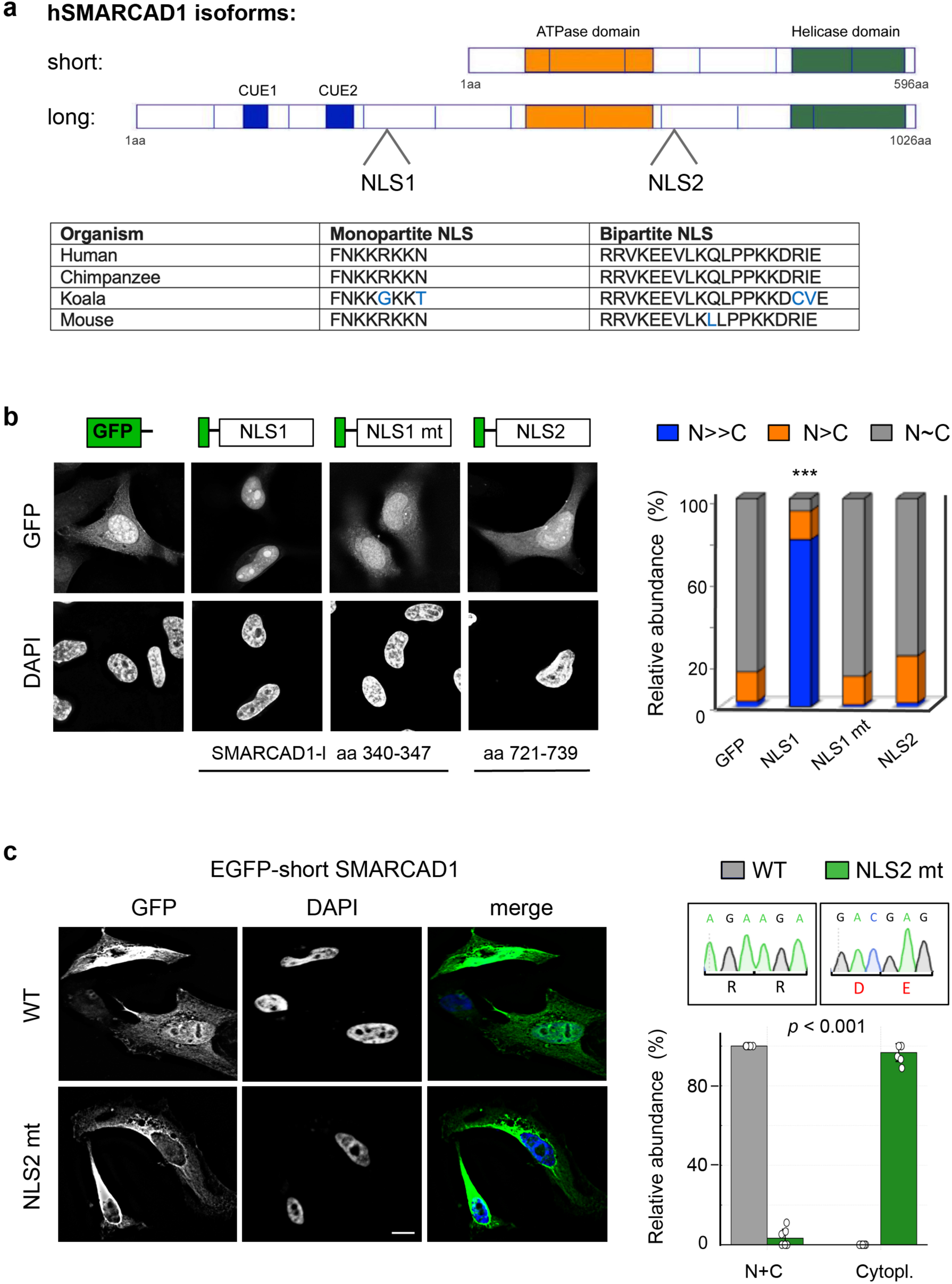
Distinct nuclear localization signals determine steady-state localization of SMARCAD1 isoforms. **(a)** Schematic of SMARCAD1 domain organization showing in silico predicted nuclear localization sequences: monopartite NLS1 (aa 340-347) and bipartite NLS2 (aa 721 to 739) with sequence conservation. **(b)** NLS1, but not NLS2, is sufficient to mediate nuclear import of a reporter. HeLa cells were transiently transfected with EGFP, EGFP-NLS1 (WT: FNKKRKKN, aa 340-347 of SMARCAD1-l; mutant: FNttReeN), or EGFP-NLS2 (RRVKEEVLKQLPPKKDRIE, aa 721-739 of SMARCAD1-). Left: representative deconvolution microscopy images. Right: percentage of cells exhibiting similar nuclear and cytoplasmic (N∼ C, grey), moderately nuclear enriched (N>C, orange), or strongly nuclear enriched (N>>C, blue) fluorescence from three independent experiments. Data are derived from scoring blinded samples (n=3), every slide contained all four samples and at least 110 green cells were counted per sample. NLS1 WT showed predominant nuclear accumulation (N>>C = 79,6 ± 6.4% SD), significantly higher than all other constructs (*p* <0.001, one way ANOVA with Tukely’s post hoc test). EGFP, NLS1 mt and NLS2 were mainly N∼C. **(c)** NLS2 mutation (R291D, R292E) abolishes SMARCAD1-s nuclear localization. Left: representative deconvolution images of HeLa cells (24 h; scale bar, 10 μm; DM5500 microscope). Scale bar: 10 μm. Right: quantification of localization of WT (grey) and NLS2 mutant (green) constructs. Bars show mean ± SD of the percentage of cells with nuclear + cytoplasmic (N+C) or cytoplasmic localization; individual data points are depicted as dots. Scored were > 135 transfected cells per condition, WT vs mt localization *p*<0.001 (unpaired two-tailed *t*-test).

To experimentally validate the in silico NLS predictions, we fused candidate NLS sequences to EGFP, a low molecular weight protein (27 kDa) that is normally dispersed throughout the cell. The NLS1 fusion protein showed a striking accumulation in the nucleus (Figure 6b, N>>C = 79,6 ± 6.4% S.D.), significantly higher than all other constructs (*p* <0.001, one way ANOVA with Tukely’s post hoc test). In contrast, EGFP-NLS2 behaved like EGFP alone, with no significant nuclear enrichment. Collectively, these results indicated that NLS1 is a strong cNLS, sufficient to mediate active import of a reporter construct, whereas NLS2 is not. Effective nuclear localization and recognition by NLS receptor generally depend on clusters of basic amino acids. Therefore, we generated an NLS1 mutant in which the basic cluster was ablated (FNKKRKKN to FNTTREEN). This mutation abolished nuclear translocation of the fusion protein (Figure 6b; N>>C = 0.9 ± 1.6% S.D.), demonstrating that the conserved lysines within the wild-type motif are essential for import. While the NLS1 motif is lacking in the short isoform (Figure 6a), we sought to test whether adding NLS1 to the C-terminus of SMARCAD1-s could alter the nucleo-cytoplamic distribution patterns described in Figure 5. The consequence was a pronounced redistribution of SMARCAD1-s protein, which became predominantly nuclear (Supplemental Figure 6c, d). Our results demonstrate that the monopartite cNLS (NLS1) is dominant over other potential targeting signals present in SMARCAD1-s.

### NLS2 is required for nuclear localization of short SMARCAD1

It remains an open question, how wild-type SMARCAD1-s is transported into the nucleus. Having ruled out a role for SMARCAD1-l in mediating nuclear localisation, we asked whether the putative bipartite NLS2 (RRVKEEVLKQLPPKKDRIE), although not sufficiently strong to direct quantitative nuclear translocation of EGFP (Figure 6b), might nevertheless mediate the nuclear localization of SMARCAD1-s. To test this, we substituted the two upstream basic residues with negatively charged amino acids (R291D und R292E) and found that the mutant SMARCAD1-s was now confined to the cytoplasm (Figure 6c, Supplemental Figure 6d). This finding established a requirement for the two arginine residues at position 291 and 292 for nuclear import in the context of the short isoform and provides an explanation for how SMARCAD1-s, which lacks NLS1, is transported into the nucleus. Importantly, these results confirm that the cellular distribution of SMARCAD1-s is regulated by an active transport mechanism.

In conclusion, we have identified two functional NLSs in SMARCAD1 which play key roles in determining the steady state localization of the individual isoforms.

## Discussion

In this study, we show that isoform-specific regulation and dosage control are key determinants of SMARCAD1-s function in health and disease. By revealing that pathogenic mutations consistently reduce SMARCAD1-s abundance, we establish reduced dosage, and thus haploinsufficiency, as the mechanistic basis of SMARCAD-syndrome. Beyond defining the disease mechanism, we delineate the regulatory processes that govern SMARCAD1-s expression and localization under physiological conditions. We show that the isoform-specific exon 1 constrains SMARCAD1-s protein output, revealing a previously unrecognized post-transcriptional regulatory layer in which upstream non-coding sequences fine-tune the abundance of a SNF2-like ATPase. We further show that SMARCAD1-s exhibits regulated nucleocytoplasmic distribution and identify NLS2 as a key determinant of its nuclear localization. Together, these findings have important implications for understanding SMARCAD1-s function in ectodermal homeostasis and provide new insights into the regulatory strategies that confer biological specificity on chromatin remodellers.

The first exon of SMARCAD1-s is vulnerable to pathogenic variation. Although most disease-associated mutations cluster at its donor splice site, it remained unclear whether they give rise to similar or distinct molecular consequences ^8,16^. Our analysis of four representative point mutations demonstrates that each disrupts pre-mRNA processing by impairing splicing of the first exon. Despite differing positions relative to the canonical GT motif, all variants caused substantial intron retention and reduced mRNA abundance in reporter assays, indicating that nucleotide changes within the +1 to +5 region compromise efficient recognition of the 5’splice site by factors involved in intron removal. Notably, one pathogenic lesion preserves the splice donor but removes sequences within and upstream of exon 1 ^11^. This 1.2 kb deletion encompasses the predicted promoter region, suggesting that disease mechanism in this case arises from impaired transcription. Together, these findings show that genetically diverse mutations converge on a common outcome, namely reduced SMARCAD1-s expression, through disruption of regulatory elements controlling either transcript production or processing.

Earlier studies proposed that SMARCAD1-s mutations cause abnormal splicing and loss of function based primarily on the c.379+1G>T substitution, which induces cryptic splice site usage ^8, 9^ . However, in our analysis, correctly spliced mRNA was detected from the +2T>C, +3A>T and +5G>C variants in skin cells. These observations indicate that SMARCAD1-s splice site mutations do not invariably result in erroneous splicing. Importantly, disruption of the exon 1 donor splice site does not preclude protein synthesis; we show that both correctly and incorrectly spliced SMARCAD1-s transcripts can be translated. As pathogenic mutations reside within upstream non-coding sequences they leave the protein sequence intact. However, they nonetheless reduce protein abundance, consistent with the observed decrease at the mRNA level. Inefficient splicing emerged as a primary contributor to reduced transcript and protein levels, as enforcing fully efficient mis-splicing restored protein output comparable to that of the wild-type. Taken together, these data suggest that patients carrying these mutations exhibit overall reduced and likely variable levels of SMARCAD1-s protein.

Clinical severity of SMARCAD-syndrome varies between families, but the basis of this variation remains unclear. We propose that differences in SMARCAD1-s levels may contribute, with relative protein concentration influencing functional outcomes. While we do not directly define a quantitative threshold, disease manifestations may arise once SMARCAD1-s levels fall below a range sufficient for normal function. Similar context-dependent dosage effects have been observed for other remodellers such as BRG1, where responses range from linear to buffered depending on the assay and the genomic locus ^63^. Notably, previously work suggested that Huriez syndrome patients exhibit lower SMARCAD1-s protein levels ^17^, but these conclusions were based on results generated with an antibody that only recognizes the long isoform. Future studies using isoform-resolved assays in patient material are therefore needed to clarify how differential SMARCAD1-s protein abundance relates to the variable clinical phenotype of ectodermal dysplasias.

Exon 1 exerts a repressive effect on SMARCAD1-s expression, with minimal impact on RNA abundance but a pronounced reduction in protein output. These findings support a model in which exon 1 modulates SMARCAD1-s protein abundance post-transcriptionally and identify upstream sequences as key contributors to SMARCAD1-s dosage control. RNA structure modelling indicates that inclusion of exon 1 reduces accessibility to the AUG start codon, potentially limiting ribosomal engagement and translational efficiency. Further support for translational control comes from the identification of multiple candidate short upstream open reading frames (uORFs) within exon 1. uORFs are well-established cis-regulatory elements that modulate translation of the main coding sequence, particularly in developmental and dosage sensitive contexts, yet their contribution to chromatin remodeller regulation has not been explored ^47,64,65^. We show that several conserved SMARCAD1-s uORFs are capable of initiating reporter translation, establishing that they are recognized by the translational machinery. This is consistent with a potential role in dampening SMARCAD1-s protein levels. While the precise mechanistic details remain to be defined, this post-transcriptional regulation operates in healthy cells and, together with evidence linking reduced SMARCAD1-s levels to disease, indicates that SMARCAD1-s abundance is bidirectionally constrained. Such tight homeostatic control is a hallmark of haploinsufficient genes, which require protein levels to be maintained within a narrow optimal range ^66^. Given the established roles of 5′ untranslated regions, uORFs and RNA secondary structure in regulating translation efficiency, our findings suggest that upstream untranslated sequences may represent an underappreciated layer of protein dosage control, particularly in dosage-sensitive genes in which precise protein abundance is essential for normal function.

Transcriptional control of SMARCAD1-s ensures tissue specific expression through alternative promoter usage (Supplemental Figure 1)^8^. Peak expression in proliferating, undifferentiated keratinocytes, followed by downregulation upon differentiation, points to a role in early epidermal development. Supporting this, perturbed expression of epidermal differentiation-associated genes has been observed in keratinocytes derived from patients carrying the c.379+1G>T SMARCAD1-s mutation ^9^, although the precise cellular function of SMARCAD1-s in skin cells remains to be fully defined. SMARCAD1-s lacks the N-terminal domain present in the long isoform, which has been implicated in targeting the remodeller to heterochromatin, replication foci and sites of DNA damage ^32,34,35,55^. This structural difference indicates that SMARCAD1-s likely performs functions distinct from the full-length protein. Moreover, although both SMARCAD1 isoforms are expressed in keratinocytes, their distinct subcellular localization suggests functional specification.

ATP-dependent chromatin remodelling enzymes are typically confined to the nucleus, with few reports describing stress-or stimulus-dependent accumulation of remodellers or co-factors in the cytoplasm ^67–69^. One example is the PBAF complex, which exhibits a cytoskeletal function during mitosis ^67^. Unlike these cases of transient relocalization, our study reveals that SMARCAD1-s is constitutively present in both the nucleus and the cytoplasm in every cell at steady-state; to our knowledge an unprecedented localization pattern for this class of enzymes. SMARCAD1-s nuclear import is an active process, dependent on a signal we designated as NLS2, ^291^RRVKEEVLKQLPPKKDRIE^309^. NLS2 has many of the characteristics of a bipartite cNLS, an upstream RR dipeptide followed by a downstream cluster containing three basic residues ^70^. Mutation of the two upstream arginines abolishes nuclear localization, confirming their functional relevance. However, the presence of two acidic amino acids (D,E) within the downstream segment could weaken its overall affinity for the receptor, importin alpha ^71^, consistent with our observation that NLS2 alone does not confer a robust nuclear enrichment when fused to GFP. We therefore infer that NLS2 functions as a comparatively weak import signal. SMARCAD1-s localization is independent of the ATPase activity, and the long isoform is not required for its compartment targeting or retention. While the observed dual distribution could reflect two distinct SMARCAD1-s subpopulations, cell-to-cell variability in nuclear versus cytoplasmic levels is suggestive of dynamic nucleo-cytoplasmic shuttling. It is conceivable that SMARCAD1 harbors a nuclear export signal (NES) that is masked in the long isoform, but becomes exposed upon N-terminal deletion. Although several candidate hydrophobic NES motifs can be identified in SMARCAD1-s, none has yet been experimentally validated.

The functional significance of the cytoplasmic localization of SMARCAD1-s remains speculative. One, SMARCAD1-s may carry out thus far unknown functions in the cytoplasm. ATP-dependent remodellers act primarily, but not exclusively, on nucleosomes ^1^. As a member of the DEAD/H box containing helicase superfamily, SMARCAD1-s could have activity towards other macromoleuclar assemblies, including RNA-protein complexes. Alternatively, it could carry out non-canonical functions beyond remodelling. Furthermore, the cytoplasmic fraction could represent an inactive reservoir, which, upon receiving appropriate cues, is transported to the nucleus. Biochemical and structural studies have revealed that in the absence of their preferred substrate, ATP-dependent chromatin remodelling enzymes are held in a self-inhibited, inactive state ^24,72^. For SMARCAD-l it has recently been demonstrated that its N-terminal domain contributes to autoinhibition ^23,24,73^. Short SMARCAD1 lacks the N-terminal domain, and hence, cytoplasmic sequestration might provide an alternative strategy for confining its activity. A cryo-EM structure of SMARCAD1-l supports the notion that once inside the nucleus, short SMARCAD1 is competent to engage chromatin, as it retains the ability to form a central DNA-binding cleft and to interact extensively with histone H4 ^24^.

Our identification of a long isoform-specific NLS (NLS1) within the N-terminus of SMARCAD1 provides a mechanistic explanation for the distinct cellular distribution of the two isoforms. This SV40-like cNLS (^340^FNKKRKKN^3^^47^) is sufficient to drive efficient nuclear import when fused to GFP or SMARCAD1-s, with activity strictly depending on an intact basic cluster. The presence of two distinct NLS motifs, one potent and isoform-specific (NLS1), the other broadly conserved but weak (NLS2), suggests that subcellular distribution of SMARCAD1 isoforms is subject to sophisticated regulation. Chromatin remodeller achieve functional diversity through multiple mechanisms, including differences in domain architecture, variable assembly with accessory factors and cell-type restricted isoforms ^2^. We propose that isoform-specific subcellular localization represents an additional layer of functional diversification in this family of remodellers.

## Methods

### Cell Culture and Transfections

HeLa cells were grown as recommended by the American Type Culture Collection (ATCC) in DMEM (Gibco) with 10% (vol/vol) FBS, 1% Penicillin-Streptomycin at 37°C and 5% CO_2_. HeLa *Smarcad1* KD cells ^32^ were additionally maintained in 2 µg/ml Puromycin. Primary epidermal keratinocytes and hTERT-KER-CT keratinocytes (KER-CT, ATCC CRL-4048) were grown in Keratinocyte Growth Medium 2 (KGM2 medium, PromoCell) with supplements including 0.06 mM CaCl_2_ at 37°C and 5% CO_2_ as described in ^74^. HaCaT keratinocytes ^50^ were grown in high Glucose DMEM (Gibco) with 10% (vol/vol) FBS, 1% Penicillin-Streptomycin, sodium pyruvate (1%) and non-essential-amino acids (1%) at 37°C and 5% CO_2_. Two fibroblast cell lines (BJ-5ta, ATCC CRL-4001, and skin fibroblasts described by ^75^ were cultured in DMEM with high glucose, supplemented with 10% FCS, 1% penicillin/streptomycin, 1% non-essential amino acids and 1% sodium pyruvate. Primary keratinocytes were isolated from breast skin patches after surgery. Human skin tissue specimens (∼0.5 cm in width) were washed in phosphate-buffered saline (PBS) and subcutaneous fat was removed. Biopsies were cut into 0.5 × 0.5 cm pieces perpendicular to the epidermis to preserve tissue architecture. Tissue fragments were incubated with Dispase (Grade I; Roche) overnight at 4 °C (or alternatively for 1–2 h at 37 °C) to separate epidermis from dermis. The epidermal layer was enzymatically dissociated using trypsin/EDTA at 37 °C for 20 min. Cells were resuspended in culture medium, filtered, washed, pooled, and seeded for monolayer growth in serum-free medium supplemented with epidermal growth factor (EGF, 0.15 ng mL⁻¹; Gibco) and bovine pituitary extract (BPE, 25 µg mL⁻¹; Gibco).

All cell lines were routinely tested with MyoAlert Detection (Lonza) for mycoplasma contamination. Transfection was carried out using Lipofectamine 2000 (HeLa and primary keratinocytes: 5 µg DNA and 8 µl Lipofectamine 2000 per 6 well) or Lipofectamine 3000 (HaCaT: 2,5 µg DNA, 3,75 µl Lipofectamine 3000 and 5 µl P3000 per 6 well) following supplier recommendations. Cells were harvested for further analysis at the times indicated at the individual figures.

### Keratinocyte Differentiation

Two independent methods of keratinocyte differentiation were used. One, differentiation was induced by cell-cell contact inhibition without addition of high calcium ^51^. Two, cells were at >90% confluency when differentiation was induced by an increase in the calcium concentration in the media through addition of 2 mM CaCl_2_ ^52^. Primary and immortalized keratinocytes were initially grown in monolayer cultures under low calcium conditions (0.05 mM CaCl_2_). hTERT-KER-CT were seeded in a density of 1.5 x 10^5^ cells per 6-well two days prior to triggering differentiation. Primary epidermal keratinocytes were seeded on 12-well plates (0.7 x 10^5^ cells per well) and grown for 3 days before inducing differentiation. For gene-expression analysis undifferentiated cells (day 0) and cells under differentiation for 96 hours (day 4) were harvested, and total RNA was extracted and analyzed by RT-qPCR.

### Protein stability assessment using MG132

V5-tagged SMARCAD1-s constructs comprising exon1, Intron1, exon2-16 were introduced into HeLa cells, and 24 hours later cells were split into two aliquots. After 1 hour, either DMSO or MG132 (20µM) was added, and after 4 hours, total protein extracts were prepared. 20 ug total protein extract was analysed and SMARCAD1-s was detected with an anti-V5-tag antibody. To assess endogenous SMARCAD1-l, Hela cells were treated for 4 hours with either DMSO or 5, 10, or 20 µM MG132.

### Plasmid construction

#### SMARCAD1 splice-site mutants

Sequence changes mimicking mutations found in ADG, Basan and Huriez syndrome patients were introduced into the 5’ splice site located at the exon1-intron1 of skin SMARCAD1 by PCR-based mutagenesis using the QuikChange II XL Site-Directed Mutagenesis Kit (Agilent 200521) according to the manufacturer’s instructions. The initial template was a minigene plasmid described by ^8^, which was used to investigate the effect of splice site variants on pre-mRNA splicing. The vector contains EGFP, human skin SMARCAD1 unique exon1, a shortened intron1 and exon2 followed by a poly A site. The intron lacks the middle 8.7 kb found in vivo, but retains the first 1.3 kb and last 0.5 kb. The WT minigene and the +1G>T mutation-contaning-plasmids were kind gifts from Eli Sprecher, Tel Aviv Sourasky Medical Center ^8^. QuikChange primers to generate additional mutations in intron1 corresponding to c.378 +2T>C, +3A>T or +5G>C were designed according to Agilent Technologies website (http://www.genomics.agilent.com), incorporating suggestions from ^76^.

To analyse the effect of splice variants on skin SMARCAD1 protein production, exon 1 and intron 1 (WT or mutations of either c.378 +1G>T or c.378 +2T>C) were cloned into an expression plasmid containing exons 2-16 of skin SMARCAD1 followed by a C-terminal V5 tag. Methodologically, this involved standard cloning techniques and PCR mutagenesis; details are available upon request. We also simulated one splice defect that was repeatedly observed when the c.378 +1G>T mutant was introduced into mammalian cells, namely a deletion of the last nucleotide of exon 1. To this end a deletion of the last G in exon 1, nucleotide 363, was introduced into the exon 1-intron 1-exons 2-16-V5 expression construct by standard cloning. The primers used to generate 5’ splice site mutations are listed in Supplemental Table 2. All plasmids were verified by sequencing.

#### Experimental validation of NLS candidate sequences

NLS candidate sequences were fused C-terminal of EGFP in the pEGFP1-C1 plasmid (Clontech) via site-directed plasmid mutagenesis PCR (Liu et al Naismith, 2008). Primers used for the one-step insertion are listed in Supplemental Table 2. The resulting DNA pool was treated with DpnI to digest the parental plasmid and transformed in NEB5alpha bacteria. Sequencing confirmed that the plasmid DNA contained the desired NLS, either WT NLS1 (FNKKRKKN corresponding to aa 340-347 of the long SMARCAD1 isoform) and NLS2 (RRVKEEVLKQLPPKKDRIE), corresponding to aa 721-739 of the long or aa 291-309 of the short SMARCAD1 isoform, or mutant NLS1 (FNTTREEN). Modification of NLS1/NLS2 within the full length SMARCAD1 isoforms is described below.

#### Mammalian expression plasmids encoding SMARCAD1 isoforms

A vector containing human short SMARCAD1 exon 1, intron 1, exons 2-16 and a C-terminal tag (V5) was prepared as follows: The main protein coding region of short SMARCAD1 (exon 2-exon 16) was PCR amplified together with a C-terminal fusion of a V5 tag from a sequence that carries silent mutations in SMARCAD1, which makes it resistant to shRNAs used in knockdown experiments (Rowbotham et al., 2011). Exon 1 and intron 1 of short SMARCAD1 were PCR amplified from the mini gene described above ^8^. Using Gibson assembly, these fragments were inserted into a pCAG (chicken beta-actin promoter)-vector, which was prepared by cutting with the restriction enzymes XhoI and NotI. Two variations of this short SMARCAD1 exon 1, intron 1, exons 2-16-V5 tag plasmid were generated, one lacks intron 1 but contains exons 1-16 and the V5 tag, and one which lacks exon 1 and instead starts with exon 2 that harbors the main ATG (short SMARCAD1 exon 2-16-V5). The first construct was generated using a gblock (IDT) and standard cloning with restriction sites, the second (short SMARCAD1 exon 2-16-V5) was created via one step PCR cloning.

Plasmids used to study the subcellular localization of SMARCAD1 exhibited the chicken beta actin promoter (pCAG). They include the short SMARCAD1 exon 1-16-V5 vector described above and plasmids that express the long and short isoforms in frame with EGFP. As the fusion of EGFP was at the N-terminus, the main ATG encoding the methionine was removed in the corresponding SMARCAD1 sequences. Moreover, silent mutations were present in the SMARCAD1 isoforms that abolish the binding of an shRNA used to generate stable SMARCAD1 knockdown (KD) HeLa cells ^32^ to prohibit the degradation of these constructs when introduced into these *Smarcad1* KD cells. Specifically, the SMARCAD1 sequence 5’-tac cag cat ttg atg acc atc aat gca -3’ replaces the original 5’-tac cag cac ctt atg aca att aat gca - 3’ sequence. The EGFP-long SMARCAD1 construct additionally displayed a V5 epitope at the N-terminus of SMARCAD1. Three variations of the WT EGFP-short SMARCAD1 plasmid exon 2-16 were generated. One, the ATPase mutation K98R, was build by changing AAA (K) to AGA (R) via one-step site directed mutagenesis. Two, the NLS1 sequence was added to skin SMARCAD1 that normally lacks this sequence. This was achieved by amplifying the skin SMARCAD1 coding region with Q5 Taq polymerase (Thermo). The PCR product was cloned into a pJET vector (Thermo) and the WT NLS1 sequence (FNKKRKKN) was inserted at the C-terminus of skin SMARCAD1 using PCR mutagenesis. Thereafter, the modified skin SMARCAD1 sequence was reintroduced in the starting vector. Primers used for the PCR mutagenesis steps are listed in Supplemental Table 2. Third, an EGFP-short SMARCAD1 plasmid was generated that carries a NLS2 mutation (R291D und R292E, corresponding to a DNA sequence change from AGAAGA to GACGAG) using a gblock (IDT) and standard cloning with restriction sites BsrGI & NotI. All plasmids were confirmed by sequencing, transfected into WT or *Smarcad1* KD HeLa cells in parallel with the WT version of SMARCAD1 and, where indicated, with a control plasmid (e.g. EGFP-macroH2A1.2), to assess the localization of the GFP and V5 fusion proteins.

### Dual luciferase reporter

DNA sequences from exon1 of short SMARCAD1 were cloned into the NheI site of the psiCHECK2 (Promega) vector. First, the native ATG start codon of hRluc was replaced with a TTG using site directed mutagenesis. The plasmid was then linearised with NheI, dephosphorylated, and the putative uORFs were inserted. These contained 6 nt upstream of their respective start codon to maintain the endogenous Kozak environment, but lacked the stop codon to be translated as a fusion protein.

### SMARCAD-s Splicing Patterns and associated Protein Expression

For splicing analysis and sequencing of the spliced products, HeLa and HaCaT cells were transfected with SMARCAD1-s minigenes comprising exon 1, intron with a WT or mutated 5’ splice site, and exon 2. After 24 or 48 hours, RNA was extracted and treated with Turbo DNase (Ambion) before reverse transcription with SuperScript II RT (Invitrogen). PCR was performed with cDNA and minus RT control using DreamTaq DNA polymerase (Thermo Scientific). SMARCAD1 sequences were amplified with primers for exon 1 and 2 using following conditions: 95°C for 3 min, followed by 30 cycles at 95°C for 30 sec, 50°C for 30 sec and 72°C for 1 min. ACTB was used as reference and amplified as follows: 95°C for 3 min, followed by 30 cycles at 95°C for 30 sec, 55°C for 30 sec and 72°C for 30 sec. The oligos used are listed in Supplemental Table 2. Untransfected cells were processed in parallel and served as a negative control. PCR products were resolved on agarose gels; the size of un-spliced transcripts and splicing product is 2226 bp and 355 bp respectively. Analysis by Sanger sequencing involved ligation of PCR products into pJET1.2/blunt (CloneJET PCR Cloning Kit, Thermo Scientific) according to the manufacturer’s instructions. Following transformation of ligation products into NEB5α cells, colony PCR using primers for SMARCAD-s was performed. Resulting amplicons were purified with QIAquick PCR purification kit (QIAGEN) and sequenced by LGC Genomics (Berlin) or Microsynth Seqlab GmbH (Göttingen). The obtained sequences were aligned and analyzed using Clustal Omega (EMBL-EBI) and SnapGene (GSL Biotech).

To examine how 5’ splice site mutations affect production of SMARCAD1-s protein, expression-plasmids harboring exon 1, intron1 with a WT or mutated 5’ splice site, exon 2-16 and a C-terminal tag (V5) were introduced into HeLa and HaCaT cells. To control for transfection efficiency across samples, cells were co-transfected with plasmids encoding an EGFP fusion protein. After 24 or 48 hours, whole cell extracts were prepared with different protocols, resolved on SDS-PAGE and immunoblotted with antibodies detecting SMARCAD1-s (V5) and the transfection control (GFP). In parallel, RNA was extracted and analysed by RT-qPCR.

### Lysate preparation, Immunopreciptiation and Western blot

Whole cell extracts were routinely prepared as described previously ^55^ using DNAse I to digest the DNA. Cells were collected by centrifugation, pellets were washed twice with ice-cold PBS with protease inhibitors, snap frozen and stored at -80°C. For protein extraction, cell pellets were quickly thawed, resuspended in a 3-5x volume of lysis buffer (20 mM HEPES, pH 7.3; 110 mM KOAc; 5 mM NaOAc; 2 mM MgOAc; 1 mM EGTA; freshly added: 2 mM DTT, 0.1 % NP-40; protease inhibitors) with 10 mM MnCl_2_ and 20 µg/ml DNase I and incubated at 37°C for 20 min ^55^. Alternatively, whole-cell extract were prepared as described by ^77^ or by cell-lysis directly in colour-less Laemmli dye (50 mM Tris HCl pH 6.8; 1% SDS; 100 mM DTT; 10% glycerol) followed by incubation in a sonification waterbath.

Protein concentration was determined by Bradford assay (Bio-Rad or Expedeon BradfordUltra). Following separation of proteins by SDS-PAGE, proteins were transferred to a nitrocellulose membrane with a semi-dry blotting system (Bio-Rad), or to a PVDF membrane by Wet-blotting. Membranes were blocked in PBS-T with 5% skimmed milk powder at room temperature for 1 hour and incubated with primary antibodies in 5 % BSA/PBS-T at 4°C overnight. Membranes were washed three times with blocking solution and incubated with HRP-coupled secondary antibodies (anti-rabbit IgG or anti-mouse IgG 1:10.000 GE Healthcare) at room temperature for 1 hour. Blots were washed again three times with PBS-T and developed using Immobilon HRP substrate solution (Millipore) and a ChemiDoc imaging system (Bio-Rad).

Primary antibodies included: SMARCAD1 HPA016737 (Sigma) 1:10.000; PAB15737 (Abnova) 1:2.500; A301-593A (Bethyl) 1:1.000; GFP (JL8, Clontech) 1:1000; GFP (11814460001 Roche) 1;2500; GAPDH [6C5] (ab8245, Abam) 1:5000; V5 (R960 Invitrogen) 1;5000; LaminB (ab16048, abcam) 1:5000. Anti-p53 (DO-1), 1;5000. Renilla luciferase (ab185925, abcam) 1;2500; Firefly luciferase (ab185924, abcam) 1;1000.

### In vitro transcription and translation

In vitro translation was carried out using the TNT^R^T7 Quick Coupled Trancription/Translation System (Promega #C8021) following the manufacturer’s protocol and employing a dual luciferase reporter vector (psiCHECK-2, Promega). The native AUG start codon of the Renilla luciferase gene was mutated to a non-start codon (TTG) using one step PCR mutagenesis, and putative upstream open reading frames from exon 1 of SMARCAD1-s were inserted at the original start site to assess their ability to initiate translation. Production of luciferase proteins was measured after 90 minutes incubation in reticulocyte lysate using Western blots.

### RNA isolation and RT-qPCR

Cells were washed 3x with medium and lysed with TRIzol reagent (Invitrogen). RNA extraction was performed according to the manufacturer’s instructions, and RNA integrity was analyzed by gel electrophoreses. Isolated RNA was treated with Turbo DNase (Ambion) and cDNA was synthesized with SuperScript II RT (Invitrogen) according to the manufacturer’s instructions. RT-qPCR was carried out using an iTaq Universal SYBR Green Supermix in a CFX Connect Real-Time PCR Detection System (Bio-Rad). Fluorescence detection and data analysis were performed with BioRad CFX Manager 2.0. Experiments were performed in triplicate with cDNA and minus RT controls and up to three references for gene expression normalization. Reference genes are indicated in the figure legends. They were chosen according to the cell type and experiment, for instance, YWHAZ and UBC were reported as suitable for differentiating keratinocytes ^78^. Primers, listed in Supplemental Table 2, were evaluated independently for their efficiency.

### Immunocytochemistry and Microscopy

Cells were seeded on positively charged glass slides and incubated at 37°C and 5% CO_2_ for approximately 2.5 hours to re-attach. Slides were rinsed with PBS before fixation and permeabilisation. GFP detection involved fixation for 5-10 min with 2% formaldehyde, followed by permeabilization in PBS-0.5% Triton X-100 for 5 min. Indirect immunofluorescence staining with an anti-SMARCAD1 antibody (Sigma HPA016737; 1:300 in blocking solution) involved fixation with 4% formaldehyde for 10 min and permeabilization with PBS-0.1% Triton X-100 for 10 min. V5 staining (Invitrogen R960-25; 1:400) was carried out after 15 min 4% formaldehyde fix and 10 min permeabilization in PBS-0.2% Triton X-100. All samples were blocked with 10% fetal calf serum in PBS at 4°C. Cells were stained with the first antibody at room temperature for 1 hour. After 3x washing with PBS, cells were incubated for 1 hour with a secondary antibody such as Alexa 488 coupled anti-IgG 1:600. Slides were washed 3x with PBS before mounting in Vectashield with DAPI. Alternatively, samples were incubated in 0.1 µg/ml DAPI/PBS for 5 minutes. After 2 washes in PBS, slides were rinsed in ddH_2_O and mounted in Mowiol (+ 25 mg/ml DABCO). Samples were analyzed, and images were acquired on either a Leica DMR fluorescence microscope or a Leica DM5500 (63 & 100x magnification). Cell lines or treatments that were directly compared (e.g. WT and KD cells) were fixed and stained simultaneously on the same slide. Representative images were collected with the same exposure times and identical post-processing, with FIJI or Adobe Photoshop CS3. Processing software: Blind deconvolution (Leica Application Suite Advanced Fluorescence, LAS AF, Version 4.0.0.11706); maximum intensity Z projection from 15-20 Z stacks (ImageJ).

<u>Pre-extraction:</u> *Smarcad1* knockdown cells ^32^ transfected with EGFP-tagged, shRNA-resistant SMARCAD1-s or EGFP-macroH2A1.2 were seeded on slides as described above. After 3 hours, the non-extracted control was rinsed with PBS and fixed with 2% formaldehyde for 5 min followed by permeabilization with 0.5% Triton-X100 for 5 min. The extraction slides were rinsed with PBS and incubated in ice-cold pre-extraction buffer (50 mM HEPES pH 7.4, 150 mM NaCl, 10 mM EGTA, 2 mM MgCl_2_, freshly added: 0.5% Triton X-100, 0.1 mM Benzamidine & 0.5 mM PMSF) for 30 - 60 seconds. Slides were rinsed with PBS and fixed with 2% formaldehyde for 5 min. Samples were mounted in Vectashield with DAPI and analyzed by fluorescence microscopy.

### Bioinformatical Analysis

CAGE (Cap Analysis of Gene Expression) data collected by the FANTOM5 consortium (https://fantom.gsc.riken.jp/5/) ^79,80^ were utilized. In silico prediction of NLS import signals was performed using "NLS mapper" ^81^, NLS "Stradamus” ^62^, PSORT-II (http://psort.nibb.ac.jp/ form2.html) ^61^ and ELM, the Eukaryotic Linear Motif resource ^82^. Splice site strength was estimated using MaxEntScan ^41^. RNA secondary structures were predicted using Vienna RNAfold 2.0 ^83–85^ using the following settings for folding and output: Minimum free energy (MFE) and partition, no isolated base pairs, RNA parameters of the Turner model (2004) and possible coaxial stacking of adjacent helices, energy rescaling to 37°C and a salt concentration of 140 mM displayed as MFE secondary structure. The protein structure was predicted with AlphaFold ^86^. The Ribo-uORF resource ^87^ was used to identify putative uORFs. All online predition programs/softwares were used with default prediction parameters.

### Statistical Analyses

Details of statistical analysis are provided in the appropriate figure legends.

### Ethical Statement

This study was approved by the Ethics Committee (81/21) of the Medical Faculty of the Phillipps-University, Marburg, which is in accordance with the tenets of the Declaration of Helsinki.

## Author contributions

MS and JEM conceived this study. MS performed splicing and differentiation experiments with the help of RT. ML generated primary keratinocytes. MS, FT, EP, PK undertook plasmid generation and analysis of the localization and expression of SMARCAD1 constructs; MS, LK conducted transfections and expression analysis. AS analysed RNA folding, JEM performed computational analysis; JEM wrote the manuscript with support from all other coauthors.

## Conflict of Interest

None.

## Acknowledgements

We thank Dr. Benedikt Buerfernt, Katrin Treutwein, Christos Kakalias and Ali Kachour for their help at earlier stages of this work. We are indebted to Dr. Lutz Zwiorek for performing surgery, to Jose Arteaga, Sophie Heidemann and Bianca Bamberger for their experimental support, to Dr. Eli Sprecher for generously sharing the mini-gene cassette, to Dr. Colin Dingwall for discussions about nuclear localization signals, and to Dr. Katrin Roth for microscopy advice. This work was supported by the Volkswagen Foundation (VolkswagenStiftung) through the initiative Experiment (97309).

## Notes

### Competing Interest Statement

The authors have declared no competing interest.

## References

1. Eustermann, S., Patel, A. B., Hopfner, K.-P., He, Y. & Korber, P. Energy-driven genome regulation by ATP-dependent chromatin remodellers. Nature reviews. Molecular cell biology 25, 309–332; 10.1038/s41580-023-00683-y (2024).

2. Gourisankar, S., Krokhotin, A., Wenderski, W. & Crabtree, G. R. Context-specific functions of chromatin remodellers in development and disease. Nature reviews. Genetics 25, 340–361; 10.1038/s41576-023-00666-x (2024).

3. Sokpor, G., Xie, Y., Rosenbusch, J. & Tuoc, T. Chromatin Remodeling BAF (SWI/SNF) Complexes in Neural Development and Disorders. Frontiers in molecular neuroscience 10, 243; 10.3389/fnmol.2017.00243 (2017).

4. Alfert, A., Moreno, N. & Kerl, K. The BAF complex in development and disease. Epigenetics & chromatin 12, 19; 10.1186/s13072-019-0264-y (2019).

5. Kadoch, C. & Crabtree, G. R. Mammalian SWI/SNF chromatin remodeling complexes and cancer: Mechanistic insights gained from human genomics. Science advances 1, e1500447; 10.1126/sciadv.1500447 (2015).

6. St Pierre, R. & Kadoch, C. Mammalian SWI/SNF complexes in cancer: emerging therapeutic opportunities. Current opinion in genetics & development 42, 56–67; 10.1016/j.gde.2017.02.004 (2017).

7. Arnaud, O., Le Loarer, F. & Tirode, F. BAFfling pathologies: Alterations of BAF complexes in cancer. Cancer letters 419, 266–279; 10.1016/j.canlet.2018.01.046 (2018).

8. Nousbeck, J. et al. A mutation in a skin-specific isoform of SMARCAD1 causes autosomal-dominant adermatoglyphia. American journal of human genetics 89, 302–307; 10.1016/j.ajhg.2011.07.004 (2011).

9. Nousbeck, J. et al. Mutations in SMARCAD1 cause autosomal dominant adermatoglyphia and perturb the expression of epidermal differentiation-associated genes. The British journal of dermatology 171, 1521–1524; 10.1111/bjd.13176 (2014).

10. Chang, X. et al. Heterozygous Deletion Impacting SMARCAD1 in the Original Kindred with Absent Dermatoglyphs and Associated Features (Baird, 1964). The Journal of pediatrics 194, 248–252.e2; 10.1016/j.jpeds.2017.11.011 (2018).

11. Loh, A. Y. T. et al. Huriez syndrome caused by a large deletion that abrogates the skin-specific isoform of SMARCAD1. The British journal of dermatology 184, 1205–1207; 10.1111/bjd.19799 (2021).

12. Loh, A. Y. T. et al. Huriez syndrome: Additional pathogenic variants supporting allelism to SMARCAD syndrome. American journal of medical genetics. Part A 188, 1752–1760; 10.1002/ajmg.a.62703 (2022).

13. Elhaji, Y. et al. Two SMARCAD1 Variants Causing Basan Syndrome in a Canadian and a Dutch Family. JID innovations : skin science from molecules to population health 1, 100022; 10.1016/j.xjidi.2021.100022 (2021).

14. Mathews, I., Wagh, S., Baby, A., Chandrashekar, L. & Dalal, A. Basan syndrome in family from South-India: A novel SMARCAD1 variant. Clinical and experimental dermatology; 10.1093/ced/llad393 (2023).

15. Valentin, M. N., Solomon, B. D., Richard, G., Ferreira, C. R. & Kirkorian, A. Y. Basan gets a new fingerprint: Mutations in the skin-specific isoform of SMARCAD1 cause ectodermal dysplasia syndromes with adermatoglyphia. American journal of medical genetics. Part A 176, 2451–2455; 10.1002/ajmg.a.40485 (2018).

16. Marks, K. C., Banks, W. R., Cunningham, D., Witman, P. M. & Herman, G. E. Analysis of two candidate genes for Basan syndrome. American journal of medical genetics. Part A **164A**, 1188–1191; 10.1002/ajmg.a.36438 (2014).

17. Günther, C. et al. SMARCAD1 Haploinsufficiency Underlies Huriez Syndrome and Associated Skin Cancer Susceptibility. The Journal of investigative dermatology 138, 1428–1431; 10.1016/j.jid.2018.01.015 (2018).

18. Lee, Y. A., Stevens, H. P., Delaporte, E., Wahn, U. & Reis, A. A gene for an autosomal dominant scleroatrophic syndrome predisposing to skin cancer (Huriez syndrome) maps to chromosome 4q23. American journal of human genetics 66, 326–330; 10.1086/302718 (2000).

19. Nieto-Benito, L. M. et al. Ectodermal dysplasia with congenital adermatoglyphia (Basan syndrome): Report of two cases presenting with extensive congenital milia. Pediatric dermatology 38, 530–532; 10.1111/pde.14512 (2021).

20. Li, M. et al. Genome-wide linkage analysis and whole-genome sequencing identify a recurrent SMARCAD1 variant in a unique Chinese family with Basan syndrome. European journal of human genetics : EJHG 24, 1367–1370; 10.1038/ejhg.2016.15 (2016).

21. Burger, B., Fuchs, D., Sprecher, E. & Itin, P. The immigration delay disease: adermatoglyphia-inherited absence of epidermal ridges. Journal of the American Academy of Dermatology 64, 974–980; 10.1016/j.jaad.2009.11.013 (2011).

22. Hamm, H., Traupe, H., Bröcker, E. B., Schubert, H. & Kolde, G. The scleroatrophic syndrome of Huriez: a cancer-prone genodermatosis. The British journal of dermatology 134, 512–518 (1996).

23. Markert, J., Zhou, K. & Luger, K. SMARCAD1 is an ATP-dependent histone octamer exchange factor with de novo nucleosome assembly activity. Science advances 7, eabk2380; 10.1126/sciadv.abk2380 (2021).

24. Hu, P. et al. Subnucleosome preference of human chromatin remodeller SMARCAD1. Nature; 10.1038/s41586-025-09100-0 (2025).

25. Schoor, M., Schuster-Gossler, K., Roopenian, D. & Gossler, A. Skeletal dysplasias, growth retardation, reduced postnatal survival, and impaired fertility in mice lacking the SNF2/SWI2 family member ETL1. Mechanisms of development 85, 73–83; 10.1016/s0925-4773(99)00090-8 (1999).

26. Sebastian-Perez, R. et al. SMARCAD1 and TOPBP1 contribute to heterochromatin maintenance at the transition from the 2C-like to the pluripotent state. eLife 12; 10.7554/eLife.87742.3 (2025).

27. Hong, F. et al. Dissecting early differentially expressed genes in a mixture of differentiating embryonic stem cells. PLoS computational biology 5, e1000607; 10.1371/journal.pcbi.1000607 (2009).

28. Sachs, P. et al. SMARCAD1 ATPase activity is required to silence endogenous retroviruses in embryonic stem cells. Nature communications 10, 1335; 10.1038/s41467-019-09078-0 (2019).

29. Navarro, C., Lyu, J., Katsori, A.-M., Caridha, R. & Elsässer, S. J. An embryonic stem cell-specific heterochromatin state promotes core histone exchange in the absence of DNA accessibility. Nature communications 11, 5095; 10.1038/s41467-020-18863-1 (2020).

30. Leeb, M., Dietmann, S., Paramor, M., Niwa, H. & Smith, A. Genetic exploration of the exit from self-renewal using haploid embryonic stem cells. Cell stem cell 14, 385–393; 10.1016/j.stem.2013.12.008 (2014).

31. Xiao, S. et al. SMARCAD1 Contributes to the Regulation of Naive Pluripotency by Interacting with Histone Citrullination. Cell reports 18, 3117–3128; 10.1016/j.celrep.2017.02.070 (2017).

32. Rowbotham, S. P. et al. Maintenance of silent chromatin through replication requires SWI/SNF-like chromatin remodeler SMARCAD1. Molecular cell 42, 285–296; 10.1016/j.molcel.2011.02.036 (2011).

33. Chakraborty, S. et al. SMARCAD1 Phosphorylation and Ubiquitination Are Required for Resection during DNA Double-Strand Break Repair. *iScience* 2, 123–135; 10.1016/j.isci.2018.03.016 (2018).

34. Lo, C. S. Y. et al. SMARCAD1-mediated active replication fork stability maintains genome integrity. Science advances 7; 10.1126/sciadv.abe7804 (2021).

35. Bantele, S. C. S. & Pfander, B. Nucleosome Remodeling by Fun30SMARCAD1 in the DNA Damage Response. Frontiers in molecular biosciences 6, 78; 10.3389/fmolb.2019.00078 (2019).

36. Sachs, P., Bergmaier, P., Treutwein, K. & Mermoud, J. E. The Conserved Chromatin Remodeler SMARCAD1 Interacts with TFIIIC and Architectural Proteins in Human and Mouse. Genes 14; 10.3390/genes14091793 (2023).

37. Uruci, S. et al. SMARCAD1 Regulates R-Loops at Active Replication Forks Linked to Cancer Mutation Hotspots (2024).

38. Tong, Z.-B., Ai, H.-S. & Li, J.-B. The Mechanism of Chromatin Remodeler SMARCAD1/Fun30 in Response to DNA Damage. Frontiers in Cell and Developmental Biology 8, 560098; 10.3389/fcell.2020.560098 (2020).

39. Alruwaili, J. & Hai, A. Adermatoglyphia. The National medical journal of India 32, 253; 10.4103/0970-258X.291296 (2019).

40. Wang, R., Helbig, I., Edmondson, A. C., Lin, L. & Xing, Y. Splicing defects in rare diseases: transcriptomics and machine learning strategies towards genetic diagnosis. Briefings in bioinformatics 24; 10.1093/bib/bbad284 (2023).

41. Yeo, G. & Burge, C. B. Maximum entropy modeling of short sequence motifs with applications to RNA splicing signals. Journal of computational biology : a journal of computational molecular cell biology 11, 377–394; 10.1089/1066527041410418 (2004).

42. García-Ruiz, S. et al. Splicing accuracy varies across human introns, tissues, age and disease. Nature communications 16, 1068; 10.1038/s41467-024-55607-x (2025).

43. Shaul, O. How introns enhance gene expression. The international journal of biochemistry & cell biology 91, 145–155; 10.1016/j.biocel.2017.06.016 (2017).

44. Pesole, G. et al. Structural and functional features of eukaryotic mRNA untranslated regions. Gene 276, 73–81; 10.1016/s0378-1119(01)00674-6 (2001).

45. Zhang, H. et al. Determinants of genome-wide distribution and evolution of uORFs in eukaryotes. Nature communications 12, 1076; 10.1038/s41467-021-21394-y (2021).

46. Calvo, S. E., Pagliarini, D. J. & Mootha, V. K. Upstream open reading frames cause widespread reduction of protein expression and are polymorphic among humans. Proceedings of the National Academy of Sciences of the United States of America 106, 7507–7512; 10.1073/pnas.0810916106 (2009).

47. Dever, T. E., Ivanov, I. P. & Hinnebusch, A. G. Translational regulation by uORFs and start codon selection stringency. Genes & development 37, 474–489; 10.1101/gad.350752.123 (2023).

48. McGillivray, P. et al. A comprehensive catalog of predicted functional upstream open reading frames in humans. Nucleic acids research 46, 3326–3338; 10.1093/nar/gky188 (2018).

49. Forrest, A. R. R. et al. A promoter-level mammalian expression atlas. Nature 507, 462–470; 10.1038/nature13182 (2014).

50. Boukamp, P. et al. Normal keratinization in a spontaneously immortalized aneuploid human keratinocyte cell line. The Journal of cell biology 106, 761–771; 10.1083/jcb.106.3.761 (1988).

51. Smits, J. P. H. et al. Immortalized N/TERT keratinocytes as an alternative cell source in 3D human epidermal models. Scientific reports 7, 11838; 10.1038/s41598-017-12041-y (2017).

52. Borowiec, A.-S., Delcourt, P., Dewailly, E. & Bidaux, G. Optimal differentiation of in vitro keratinocytes requires multifactorial external control. PloS one 8, e77507; 10.1371/journal.pone.0077507 (2013).

53. Glover, J. D. et al. The developmental basis of fingerprint pattern formation and variation. Cell 186, 940–956.e20; 10.1016/j.cell.2023.01.015 (2023).

54. Okazaki, N. et al. The novel protein complex with SMARCAD1/KIAA1122 binds to the vicinity of TSS. Journal of molecular biology 382, 257–265; 10.1016/j.jmb.2008.07.031 (2008).

55. Ding, D. et al. The CUE1 domain of the SNF2-like chromatin remodeler SMARCAD1 mediates its association with KRAB-associated protein 1 (KAP1) and KAP1 target genes. The Journal of biological chemistry 293, 2711–2724; 10.1074/jbc.RA117.000959 (2018).

56. Mermoud, J. E., Rowbotham, S. P. & Varga-Weisz, P. D. Keeping chromatin quiet: how nucleosome remodeling restores heterochromatin after replication. *Cell cycle (Georgetown*, Tex.) 10, 4017–4025; 10.4161/cc.10.23.18558 (2011).

57. Awad, S., Ryan, D., Prochasson, P., Owen-Hughes, T. & Hassan, A. H. The Snf2 homolog Fun30 acts as a homodimeric ATP-dependent chromatin-remodeling enzyme. The Journal of biological chemistry 285, 9477–9484; 10.1074/jbc.M109.082149 (2010).

58. Adra, C. N. et al. SMARCAD1, a novel human helicase family-defining member associated with genetic instability: cloning, expression, and mapping to 4q22-q23, a band rich in breakpoints and deletion mutants involved in several human diseases. Genomics 69, 162–173; 10.1006/geno.2000.6281 (2000).

59. Lu, J. et al. Types of nuclear localization signals and mechanisms of protein import into the nucleus. Cell communication and signaling : CCS 19, 60; 10.1186/s12964-021-00741-y (2021).

60. Tessier, T. M., MacNeil, K. M. & Mymryk, J. S. Piggybacking on Classical Import and Other Non-Classical Mechanisms of Nuclear Import Appear Highly Prevalent within the Human Proteome. Biology 9; 10.3390/biology9080188 (2020).

61. Horton, P. et al. WoLF PSORT: protein localization predictor. Nucleic acids research 35, W585–7; 10.1093/nar/gkm259 (2007).

62. Nguyen Ba, A. N., Pogoutse, A., Provart, N. & Moses, A. M. NLStradamus: a simple Hidden Markov Model for nuclear localization signal prediction. BMC bioinformatics 10, 202; 10.1186/1471-2105-10-202 (2009).

63. Hagihara, Y., Zhang, C. & Zhang, Y. Precise modulation of BRG1 levels reveals features of mSWI/SNF dosage sensitivity. Nature genetics 57, 2250–2263; 10.1038/s41588-025-02305-z (2025).

64. Wieder, N. et al. Differences in 5’untranslated regions highlight the importance of translational regulation of dosage sensitive genes. Genome biology 25, 111; 10.1186/s13059-024-03248-0 (2024).

65. Kong, J. & Lasko, P. Translational control in cellular and developmental processes. Nature reviews. Genetics 13, 383–394; 10.1038/nrg3184 (2012).

66. Morrill, S. A. & Amon, A. Why haploinsufficiency persists. Proceedings of the National Academy of Sciences of the United States of America 116, 11866–11871; 10.1073/pnas.1900437116 (2019).

67. Karki, M. et al. A cytoskeletal function for PBRM1 reading methylated microtubules. Science advances 7; 10.1126/sciadv.abf2866 (2021).

68. Lee, J. H. et al. Cytoplasmic localization and nucleo-cytoplasmic shuttling of BAF53, a component of chromatin-modifying complexes. Molecules and cells 16, 78–83 (2003).

69. Dastidar, R. G. et al. The nuclear localization of SWI/SNF proteins is subjected to oxygen regulation. Cell & bioscience 2, 30; 10.1186/2045-3701-2-30 (2012).

70. Dingwall, C. & Laskey, R. A. Nuclear targeting sequences--a consensus? Trends in biochemical sciences 16, 478–481; 10.1016/0968-0004(91)90184-w (1991).

71. Conti, E., Uy, M., Leighton, L., Blobel, G. & Kuriyan, J. Crystallographic analysis of the recognition of a nuclear localization signal by the nuclear import factor karyopherin alpha. Cell 94, 193–204; 10.1016/s0092-8674(00)81419-1 (1998).

72. Blessing, C., Knobloch, G. & Ladurner, A. G. Restraining and unleashing chromatin remodelers - structural information guides chromatin plasticity. Current opinion in structural biology 65, 130–138; 10.1016/j.sbi.2020.06.008 (2020).

73. Aboulache, B. L., Hoitsma, N. M. & Luger, K. Phosphorylation regulates the chromatin remodeler SMARCAD1 in nucleosome binding, ATP hydrolysis, and histone exchange. The Journal of biological chemistry, 107893; 10.1016/j.jbc.2024.107893 (2024).

74. Beckert, B. et al. Immortalized Human hTert/KER-CT Keratinocytes a Model System for Research on Desmosomal Adhesion and Pathogenesis of Pemphigus Vulgaris. International journal of molecular sciences 20; 10.3390/ijms20133113 (2019).

75. Mussche, S. et al. Restoration of cytoskeleton homeostasis after gigaxonin gene transfer for giant axonal neuropathy. Human gene therapy 24, 209–219; 10.1089/hum.2012.107 (2013).

76. Zheng, L., Baumann, U. & Reymond, J.-L. An efficient one-step site-directed and site-saturation mutagenesis protocol. Nucleic acids research 32, e115; 10.1093/nar/gnh110 (2004).

77. Thompson, P. J. et al. hnRNP K coordinates transcriptional silencing by SETDB1 in embryonic stem cells. PLoS genetics 11, e1004933; 10.1371/journal.pgen.1004933 (2015).

78. Lanzafame, M. et al. Reference genes for gene expression analysis in proliferating and differentiating human keratinocytes. Experimental dermatology 24, 314–316; 10.1111/exd.12657 (2015).

79. Lizio, M. et al. Gateways to the FANTOM5 promoter level mammalian expression atlas. Genome biology 16, 22; 10.1186/s13059-014-0560-6 (2015).

80. Abugessaisa, I. et al. FANTOM enters 20th year: expansion of transcriptomic atlases and functional annotation of non-coding RNAs. Nucleic acids research 49, D892–D898; 10.1093/nar/gkaa1054 (2021).

81. Kosugi, S., Hasebe, M., Tomita, M. & Yanagawa, H. Systematic identification of cell cycle-dependent yeast nucleocytoplasmic shuttling proteins by prediction of composite motifs. Proceedings of the National Academy of Sciences of the United States of America 106, 10171–10176; 10.1073/pnas.0900604106 (2009).

82. Kumar, M. et al. The Eukaryotic Linear Motif resource: 2022 release. Nucleic acids research 50, D497–D508; 10.1093/nar/gkab975 (2022).

83. Hofacker, I. L. Vienna RNA secondary structure server. Nucleic acids research 31, 3429–3431; 10.1093/nar/gkg599 (2003).

84. Lorenz, R. et al. ViennaRNA Package 2.0. Algorithms for molecular biology : AMB 6, 26; 10.1186/1748-7188-6-26 (2011).

85. Gruber, A. R., Lorenz, R., Bernhart, S. H., Neuböck, R. & Hofacker, I. L. The Vienna RNA websuite. Nucleic acids research 36, W70–4; 10.1093/nar/gkn188 (2008).

86. Varadi, M. et al. AlphaFold Protein Structure Database: massively expanding the structural coverage of protein-sequence space with high-accuracy models. Nucleic acids research 50, D439–D444; 10.1093/nar/gkab1061 (2022).

87. Liu, Q. et al. Ribo-uORF: a comprehensive data resource of upstream open reading frames (uORFs) based on ribosome profiling. Nucleic acids research 51, D248–D261; 10.1093/nar/gkac1094 (2023).

